# The ArcZ small RNA activates SlyA production to control contact-dependent antibacterial killing and diffusible antifungal activity in *Dickeya solani*

**DOI:** 10.64898/2026.09.28.754996

**Authors:** Quentin Dubois, Marcel Sprenger, Sebastian Krautwurst, Agnès Rodrigue, Kai Papenfort, Laetitia Attaiech, Erwan Gueguen

## Abstract

Bacterial phytopathogens deploy diverse antimicrobial systems, including diffusible secondary metabolites and contact-dependent secretion apparatuses, to compete with coinhabiting microorganisms during host colonization. How the production of these distinct weapons is coordinated remains incompletely understood. Here, we identified a two-tier regulatory cascade by which the Hfq-dependent small RNA (sRNA) ArcZ controls both antibacterial and antifungal antagonism in the phytopathogen *Dickeya solani*. High-throughput RNA interactome analysis (RIL-seq) shows that ArcZ interacts broadly across the transcriptome and directly base-pairs with the 5′ untranslated region of the mRNA encoding the MarR-family transcription factor SlyA, activating its translation. SlyA subsequently binds directly to selected promoter regions within the solanimycin biosynthetic cluster (*sol*) and the type VI secretion system (T6SS) loci and activates their transcription. The ArcZ-SlyA axis is required for solanimycin-mediated antifungal activity against *Kluyveromyces lactis* and for T6SS-dependent killing of the sympatric pathogen *Pectobacterium atrosepticum*, demonstrating that the *D. solani* T6SS mediates interbacterial competition. A compensatory base-pairing mutation between ArcZ and *slyA* restores both competitive phenotypes, establishing this post-transcriptional interaction as a direct upstream determinant. Together, these results reveal an sRNA-controlled regulatory axis that coordinates diffusible antifungal and contact-dependent antibacterial activities.

**Importance:** To colonize host tissues successfully, bacteria deploy specialized weapons against diverse competitors, including rival bacteria and fungi. How bacteria coordinate distinct competitive systems within a unified regulatory program remains poorly resolved. This study demonstrates that the small RNA ArcZ expands its regulatory range by activating the transcription factor SlyA, rather than targeting downstream effector genes individually. This hierarchical architecture enables a single post-transcriptional checkpoint to control both a diffusible antifungal metabolite and a contact-dependent antibacterial secretion system. Although ArcZ and SlyA are broadly conserved across *Enterobacterales*, the complementary sequence within the *slyA* 5′ UTR is restricted to *Pectobacteriaceae*, suggesting that this regulatory interaction emerged through target-site evolution. These findings provide a model for how the evolution of a non-coding target site can recruit conserved regulatory components into a lineage-specific regulatory circuit.

## Introduction

Bacterial phytopathogens of the family *Pectobacteriaceae*, particularly species of *Dickeya* and *Pectobacterium*, cause substantial agricultural losses through the maceration of plant tissues (1, 2). Although disease development depends largely on the coordinated secretion of plant cell wall-degrading enzymes (PCWDEs), successful host colonization also requires direct competition with coinhabiting microorganisms in the infected tissue. *Pectobacteriaceae* therefore deploy diverse antagonistic mechanisms, ranging from diffusible antimicrobial metabolites, produced by non-ribosomal peptide synthetase (NRPS) and polyketide synthase (PKS) pathways, to contact-dependent killing systems such as the type VI secretion system (T6SS) (3).

The emerging potato pathogen *Dickeya solani* (4, 5) possesses a broad repertoire of competitive systems. These include biosynthetic gene clusters for the antifungal macrolide oocydin A (*ooc*), the antibacterial polyamino-amide zeamine (*zms*), and the antifungal compound solanimycin (*sol*), alongside a functional T6SS (6–10). Previous work identified the small regulatory RNA (sRNA) ArcZ as an important regulator of antimicrobial activity in *D. solani*, including the expression of the *sol* and *zms* clusters (10). ArcZ is an Hfq-dependent sRNA conserved across *Enterobacterales* that controls diverse physiological processes through base pairing with target mRNAs (11). It is transcribed as a ∼132-nt precursor and processed by RNase E into a stable 59-nt isoform that constitutes its active regulatory form (12). However, bioinformatic analyses failed to detect direct base pairing between ArcZ and transcripts within the *sol* locus, suggesting that ArcZ controls solanimycin production through an intermediate regulatory factor.

A candidate regulator linking ArcZ to downstream antimicrobial systems is the transcription factor SlyA. SlyA belongs to the MarR family of dimeric winged helix-turn-helix DNA-binding proteins and is widely conserved across *Enterobacterales* (13, 14). Members of the SlyA/RovA subfamily regulate virulence, stress adaptation, and microbial antagonism, acting through direct promoter regulation or, at some loci, by counteracting H-NS-mediated transcriptional silencing (14, 15). In *Yersinia pseudotuberculosis*, the SlyA/RovA-family regulator RovA directly binds the T6SS promoter and positively regulates its expression (16). In *Dickeya dadantii* 3937, SlyA contributes to virulence, environmental stress survival, and type III secretion system expression (17, 18). In *Dickeya zeae* EC1, SlyA activates its own transcription as well as the *zms* cluster (19), integrating upstream signals from quorum sensing, OhrR, ArcAB, and integration host factor (20–22). Transcriptomic analyses have further linked SlyA to T6SS gene expression in *Pectobacterium brasiliense* and *D. zeae* (19, 23). Although SlyA functions as a central transcriptional hub in *Pectobacteriaceae*, whether its expression is subject to post-transcriptional control by sRNAs has not been determined.

Among the antagonistic systems in *D. solani*, the T6SS represents a potentially major, yet uncharacterized, contact-dependent competitive weapon. Comparative genomics has indicated a distinct repertoire of T6SS effector-immunity modules in *D. solani* (24), but its functional role in interbacterial antagonism has remained unverified, in contrast to related enterobacterial models (25, 26). Furthermore, how *D. solani* coordinates contact-dependent mechanical weapons with diffusible chemical metabolites within a unified regulatory circuit remains unknown.

In this study, we focused on ArcZ-dependent antimicrobial functions and demonstrate that SlyA provides the direct regulatory link connecting ArcZ to intermicrobial competition in *D. solani*. Using RIL-seq, quantitative proteomics and allele-specific compensatory mutations, we show that ArcZ directly base-pairs with the 5′ untranslated region (UTR) of *slyA* mRNA to promote its translation. In turn, SlyA induces transcription from selected *sol* and T6SS promoters *in vivo* and directly binds these promoter regions in *vitro*. This regulatory axis governs both solanimycin-dependent antifungal activity against *Kluyveromyces lactis* and T6SS-dependent killing of the phytopathogen *Pectobacterium atrosepticum*. A compensatory mutation that re-establishes ArcZ-*slyA* base pairing rescues both competitive outputs, confirming that this sRNA-mRNA interaction acts as a causal upstream determinant. These findings identify a multilayered regulatory architecture through which an sRNA coordinates diffusible and contact-dependent antimicrobial weapons.

## Results

### Proteomic profiling links ArcZ to the T6SS, solanimycin, and global regulators

All experiments were performed in *D. solani* D s0432-1, which carries the functional *arcZ_1_* allele and in which ArcZ was previously shown to be required for the production of the antifungal secondary metabolite solanimycin (10). Subsequent quantitative proteomic profiling of a Δ*arcZ* mutant was focused on classical plant-maceration traits, identifying ArcZ as a positive regulator of plant cell wall-degrading enzymes and 2,3-butanediol fermentation (27). However, whether ArcZ controls other competitive weapons and regulatory networks remained unexplored.

Here, re-examination of this proteomic dataset (27) showed that ArcZ deficiency leads to a marked, simultaneous reduction of the contact-dependent type VI secretion system (T6SS), an observation not reported in our previous study. Core structural and effector components, including the contractile sheath subunit TssC (log_2_FC = -1.10), the spike protein VgrGA (log_2_FC = -1.29), and the Rhs immunity protein RhsIA (log_2_FC = -1.36) were significantly reduced in Δ*arcZ* cells relative to the wild-type strain. We also observed a significantly reduced abundance of proteins involved in the biosynthesis of solanimycin, a PKS/NRPS antifungal metabolite. Levels of SolD, SolH, SolF and SolG were significantly reduced in the Δ*arcZ* cells relative to the wild-type strain, as previously described (27) (Fig. 1A). Because bioinformatic predictions failed to detect direct base-pairing interactions between ArcZ and transcripts within the *sol* cluster or T6SS operons, we hypothesized an additional level of regulation within this axis. Searching for intermediate regulators within the Δ*arcZ* proteome revealed alterations across multiple global transcription factors: the MarR-family regulator SlyA (log_2_FC = -1.62) and the alternative σ factor RpoS (log_2_FC = -1.16) were significantly depleted, whereas the global virulence repressor PecT was strongly enriched (log_2_FC = +1.6) (Fig. 1A). However, because global proteomic profiling reflects a complex mixture of direct transcriptional and post-transcriptional events, and downstream indirect consequences, these data alone could not decipher whether SlyA, PecT, RpoS, or an uncharacterized factor acts as the primary upstream driver coordinating T6SS assembly and solanimycin biosynthesis.

### RIL-seq identifies *slyA* mRNA as the dominant ArcZ mRNA target at low cell density in *D. solani*

To identify direct ArcZ targets that could mediate its effects on antimicrobial gene expression, we performed Hfq RIL-seq at low cell density (LCD; OD_600_ = 0.5; Fig. 1B) and high cell density (HCD; OD_600_ = 1.0; Fig. S1A). Northern blot analysis confirmed that ArcZ accumulates predominantly as its biologically active, processed ∼59-nt isoform during growth (Fig. S1E). Across both growth phases, ArcZ established more distinct RNA-RNA interactions than any other Hfq-associated sRNA (Fig. S1B and C, Table S1), identifying ArcZ as the most connected Hfq-dependent sRNA in *D. solani* under these conditions. No ArcZ chimeric reads were detected with transcripts within the *sol* biosynthetic cluster or within the core T6SS loci (including the primary *tss* apparatus and *hcp* tube operons). Instead, at LCD, the mRNA encoding the transcription factor SlyA emerged as the single most abundant ArcZ partner (Fig. 1C, Table S1), accounting for approximately 25% of all recovered ArcZ-mRNA chimeric reads (∼400 chimeric reads recovered consistently across all three biological replicates; Fig. S1D). Furthermore, RIL-seq ligation junctions mapped predominantly to the 5′ untranslated region (5′ UTR) of *slyA*, identifying the *slyA* leader as a prominent direct ArcZ interaction site and *slyA* mRNA as the dominant ArcZ target at low cell density.

**Figure 1.**
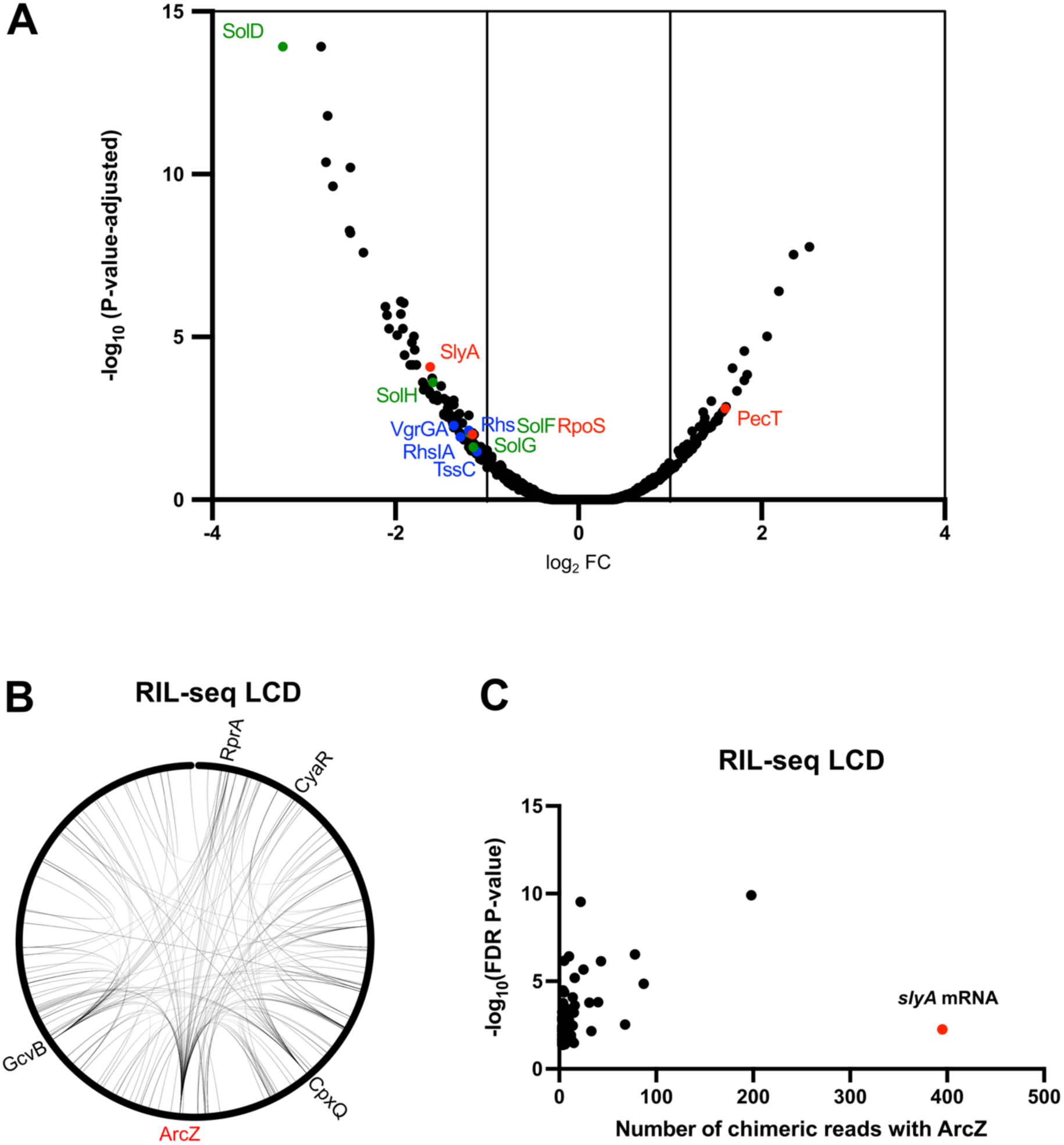
Proteomic and RIL-seq analyses link ArcZ to antimicrobial systems and identify *slyA* mRNA as its major direct target. (A) Volcano plot of quantitative proteomic analysis comparing Δ*arcZ* and wild-type *D. solani* cells grown in M63 minimal medium supplemented with 1% sucrose at 30°C to OD_600_ = 0.3. The x-axis shows the log_2_ protein abundance ratio (Δ*arcZ*/WT), and the y-axis shows -log_10_ P value. Differentially abundant proteins were defined with log_2_ FC ≤-1 or ≥ 1 and p-value ≤ 0.05. Solanimycin-associated proteins, T6SS-associated proteins, and global regulators are highlighted in green, blue, and red, respectively. Selected proteins are labeled. Data are from three independent biological replicates. The dataset was generated in ref. 27 and is reanalyzed here; the complete proteome is available in that study. (B) Hfq-dependent RNA-RNA interaction network identified by RIL-seq in *D. solani* cells grown in LB at 30°C to low cell density (LCD; OD_600_ = 0.5). Nodes represent Hfq-associated RNAs and connecting lines represent RNA pairs supported by ≥ 5 chimeric reads and an FDR base-pairing ≤ 0.05. ArcZ is highlighted in red. The circle around it represents the *D. solani* chromosome, showing the positions of the corresponding sRNAs. Data are from 3 independent biological replicates (Table S1). (C) ArcZ-interacting mRNAs identified by RIL-seq at LCD. Each point represents an ArcZ-mRNA interaction supported by ≥ 5 chimeric reads and an FDR base-pairing ≤ 0.05; *slyA* is highlighted in red. Approximately 400 ArcZ-*slyA* chimeric reads were recovered under this condition.

**Figure S1.**
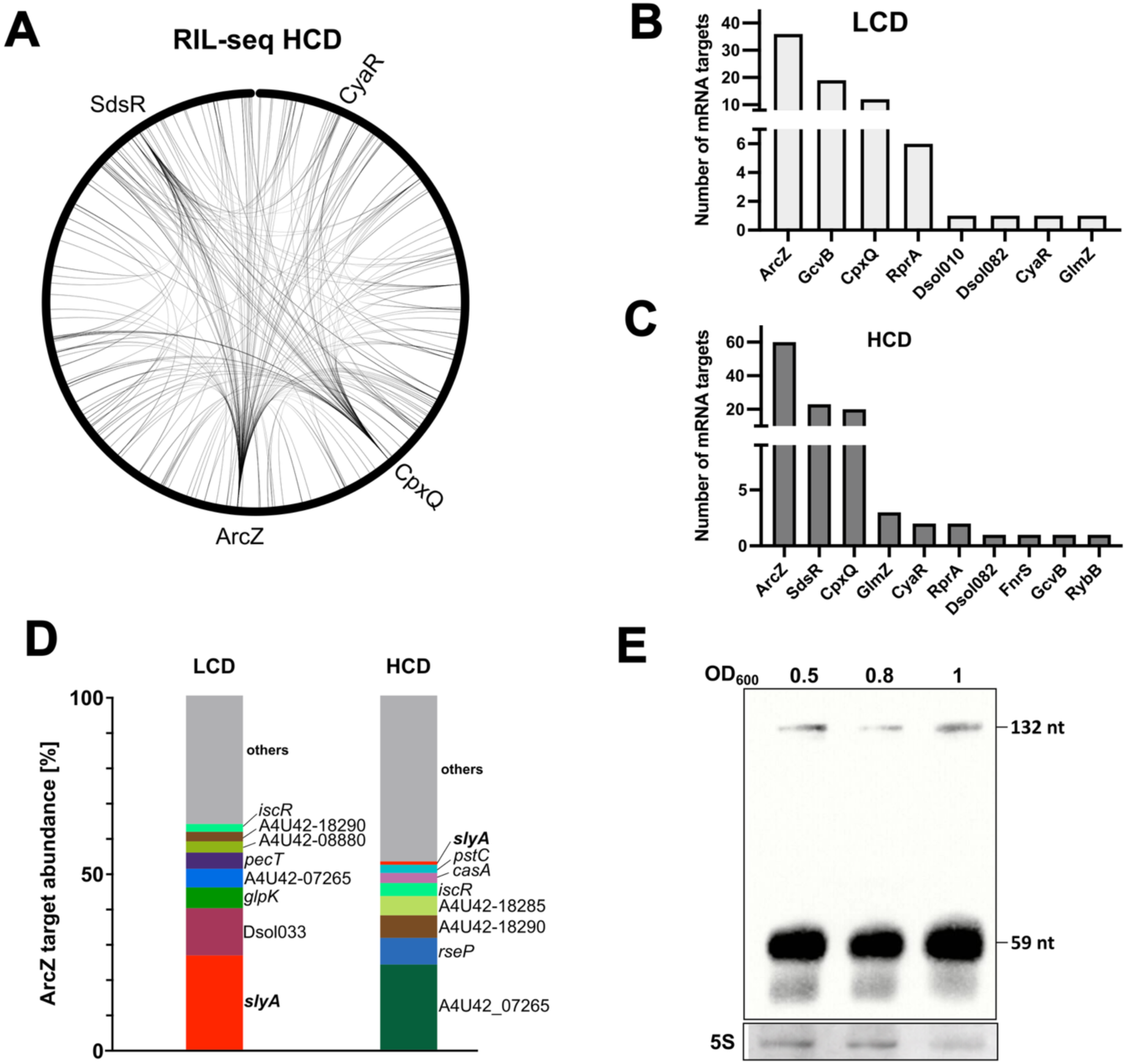
ArcZ is the most highly connected Hfq-associated sRNA and accumulates predominantly as a processed isoform. (A) Hfq-dependent RNA-RNA interaction network identified by RIL-seq in *D. solani* cells at high cell density (HCD). Connecting lines represent RNA pairs supported by ≥ 5 chimeric reads and an FDR base-pairing ≤ 0.05. Selected Hfq-associated sRNAs are indicated around the network. The circle around it represents the *D. solani* chromosome, showing the positions of the corresponding sRNAs. (B-C) Number of distinct mRNA targets identified for individual Hfq-associated sRNAs at low cell density (LCD; B) and high cell density (HCD; C). Only interactions supported by ≥ 5 chimeric reads and an FDR base-pairing ≤ 0.05 were included. Bars are shown in white for LCD and grey for HCD. (D) Relative abundance of ArcZ-interacting mRNAs at LCD and HCD. The most abundant ArcZ targets are shown individually and labeled; all remaining targets are grouped as “others” in grey. Target abundance is expressed as the percentage of total ArcZ-mRNA chimeric reads recovered under each condition. (E) Northern blot analysis of ArcZ accumulation at OD_600_ = 0.5, 0.8, and 1. The full-length ArcZ precursor (∼132 nt) and processed ∼59-nt isoform are indicated. 5S rRNA served as a loading control. A representative blot from three independent biological replicates is shown.

### ArcZ promotes *slyA* mRNA translation via direct base pairing with its 5′ UTR

To determine how ArcZ regulates *slyA* expression, we used a chromosomal translational reporter integrated at the native *slyA* locus. In this construct, the native *slyA* promoter and 5′ UTR drive expression of a *slyA*-sfGFP translational fusion, allowing ArcZ-dependent changes in SlyA translation to be monitored directly. Deletion of *arcZ* reduced the activity of the translational *slyA*-*sfGFP* fusion by approximately 5-fold, and this defect was rescued by ectopic expression of *arcZ in trans* (Fig. 2A, left), while bacterial growth kinetics remained comparable across all strains (Fig. 2A, right). Consistent with the reporter measurements, quantitative proteomic profiling (27) showed a significant 3.1-fold reduction in SlyA abundance in Δ*arcZ* cells relative to wild type (log_2_FC = -1.62) (Fig. 1A). Together, the reporter and proteomic data show that ArcZ is required for SlyA accumulation.

RIL-seq chimeras combined with IntaRNA predictions identified a complementary base-pairing region between the processed 3′ region of ArcZ (nucleotide positions 75 to 88 of the full-length ArcZ precursor) and the *slyA* mRNA 5′ UTR (Fig. 2B). To experimentally define the *slyA* 5′ UTR, we mapped its transcription start site by 5′ RACE. The transcription start site was located 94 nt upstream of the *slyA* start codon, defining a 5′ UTR that encompasses the predicted ArcZ-binding site (Fig. S2A). To verify whether this predicted duplex mediates regulation *in vivo*, point mutations were introduced into the interaction site. Nucleotide substitutions within the *slyA* mRNA 5′ UTR target site (*slyA*\*) strongly reduced translational reporter activity, and expression of wild-type ArcZ failed to restore it. Conversely, co-expression of an ArcZ variant carrying compensatory substitutions (ArcZ*) restored expression of the *slyA*\* translational reporter to levels comparable to the wild-type construct (Fig. 2C). Growth kinetics were comparable across all strains (Fig. S2B), indicating that the observed reporter differences were not attributable to growth defects. This compensatory genetic rescue demonstrates that ArcZ directly activates *slyA* mRNA translation via base pairing with its 5′ UTR.

We next evaluated whether ArcZ binding alters the accessibility of the *slyA* RBS. Secondary-structure predictions by RNAfold suggested that the RBS is sequestered within a stem-loop in the unbound *slyA* mRNA 5′ UTR (Fig. 2D, left). Formation of the ArcZ-*slyA* mRNA duplex is predicted to resolve this secondary structure, exposing the RBS to the translation machinery (Fig. 2D, right). This structural rearrangement is also supported by three-dimensional AlphaFold3 models, which predict greater exposure of the *slyA* RBS upon ArcZ binding (Fig. S2C). These models support a mechanism in which ArcZ promotes translation initiation by relieving sequestration of the *slyA* RBS.

Finally, we analyzed the phylogenetic conservation of the ArcZ-*slyA* mRNA interaction across *Enterobacterales*. The ArcZ sequence involved in the pairing with *slyA* mRNA is broadly conserved across the examined *Enterobacterales* species, whereas the complementary motif in the *slyA* mRNA 5′ UTR is restricted to members of the *Pectobacteriaceae* (Fig. 2E; Fig. S3). *In silico* binding calculations predicted thermodynamically favorable ArcZ-*slyA* mRNA interactions across *Pectobacteriaceae*, but no comparable stable interaction was predicted in representative non-*Pectobacteriaceae* taxa (Fig. S3A). Sequence alignment confirmed the conservation of the *slyA* target motif specifically within *Pectobacteriaceae* (Fig. 2E; Fig. S3B). These analyses indicate that sequence divergence within the *slyA* mRNA 5′ UTR target site, rather than variation in the ArcZ sRNA itself, determines the lineage-specific distribution of this regulatory circuit.

**Figure 2.**
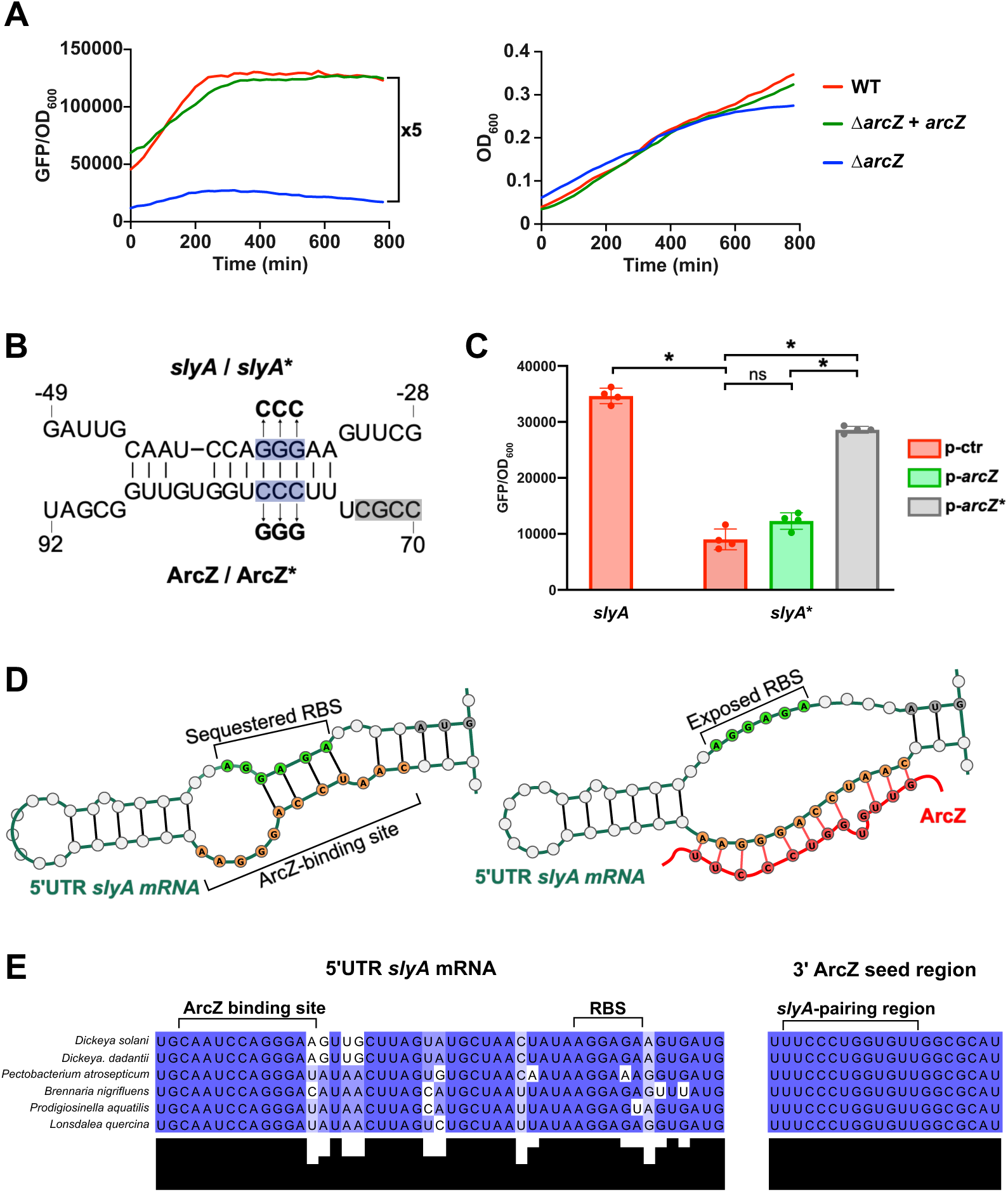
ArcZ directly activates *slyA* translation through base pairing with its 5′ UTR. (A) Effect of ArcZ on *slyA* expression. Left, activity of the translational *slyA*-sfGFP fusion, monitored as GFP fluorescence normalized to OD_600_ using a Tecan microplate reader. Right, growth of the corresponding strains monitored by OD_600_. Curves correspond to wild type (WT, red), Δ*arcZ* complemented with *arcZ* in trans (green), and Δ*arcZ* (blue). (B) Predicted base-pairing region between the processed 3′ region of ArcZ and the *slyA* 5′ UTR based on RIL-seq chimeras (28) and IntaRNA prediction. The *slyA* nucleotide substitutions introduced into *slyA* (*slyA*\*) and *arcZ* (*arcZ*\*) are indicated in bold above the original sequence. (C) Activity of the translational *slyA*-sfGFP reporter carrying the wild-type or *slyA*\* 5′ UTR in cells containing the empty control plasmid (p-ctr, red), expressing wild-type ArcZ (p-*arcZ*, green), or expressing the compensatory ArcZ* variant (p-*arcZ*\*, grey). GFP fluorescence was normalized to OD_600_, and values shown correspond to measurements collected at the end of exponential growth for 4 biological replicates. (D) Graphical representation of predicted secondary structures of the *slyA* 5′ UTR in the absence (left) and presence (right) of ArcZ. Structures were predicted using RNAfold and RNAcofold, respectively, and visualized using VARNA. The *slyA* 5′ UTR is shown in dark green, the RBS in bright green, the ArcZ-binding region within *slyA* is highlighted in orange, and ArcZ is shown in red. (E) Sequence alignment of the *slyA* mRNA 5′ UTR and the ArcZ region involved in *slyA* pairing from the indicated *Pectobacteriaceae*. The ArcZ-binding site and RBS within the *slyA* 5′ UTR and the corresponding slyA-pairing region of ArcZ are indicated above the alignments. Blue shading indicates nucleotide conservation. Complete sequence alignments and predicted ArcZ-*slyA* interactions are provided in Fig. S3. (A and C) Reporter experiments were performed with four independent biological replicates. Data are presented as mean ± SD. Statistical comparisons were performed using pairwise Mann Whitney test; *P < 0.05; ns, not significant.

**Figure S2.**
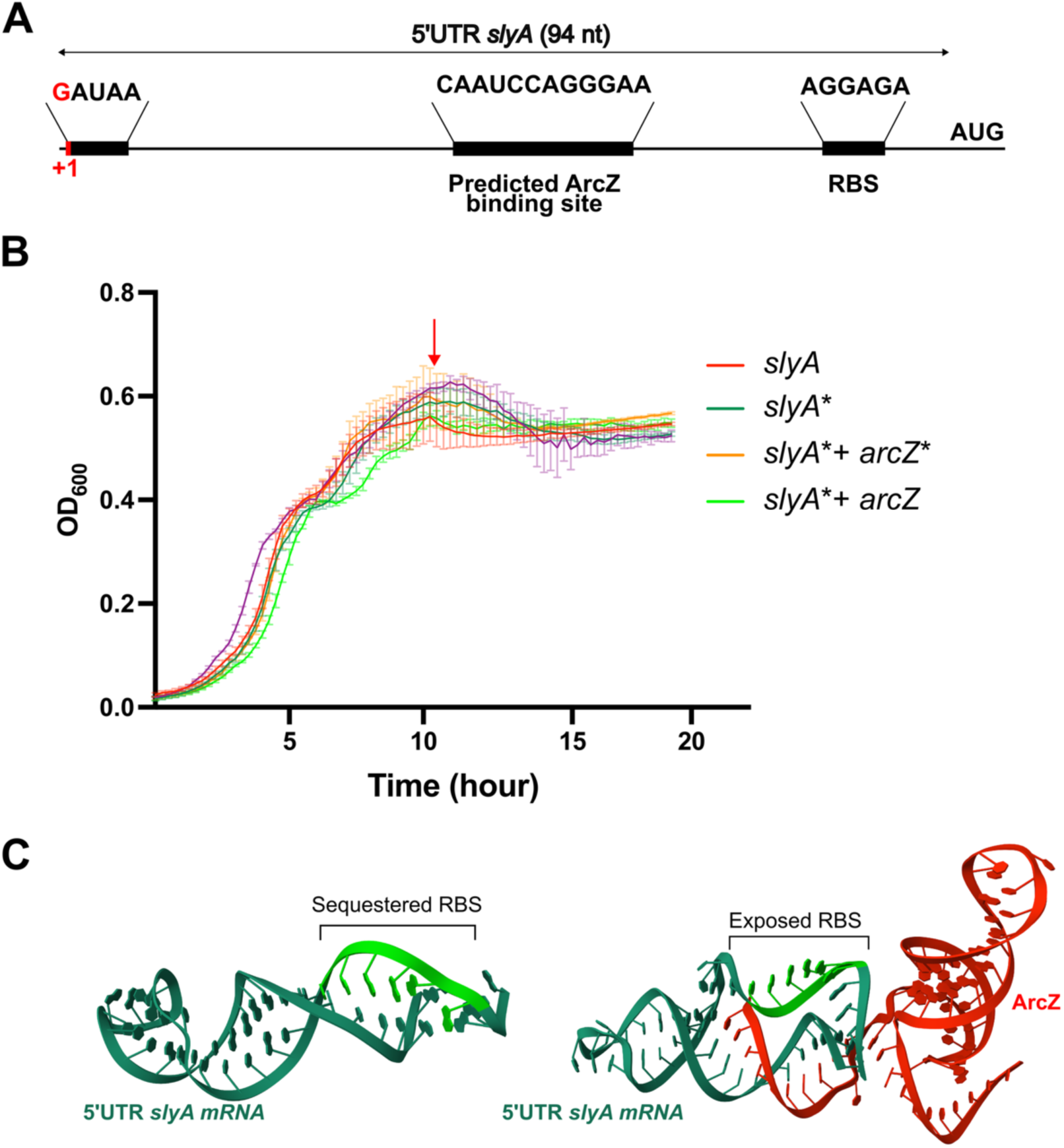
Mapping of the *slyA* 5′ UTR and structural prediction of the ArcZ-*slyA* mRNA duplex. (A) Mapping of the *slyA* 5′ untranslated region. The transcription start site TSS (+1), identified by 5′ RACE, defines a 94-nt 5′ UTR upstream of the *slyA* start codon. The predicted ArcZ-binding site, ribosome-binding site (RBS), and AUG start codon are indicated. Selected nucleotide sequences corresponding to the TSS region, ArcZ-binding site, and RBS are shown above the schematic. (B) Growth curves of the strains used in Fig. 2C, monitored by OD_600_. Curves correspond to *slyA* (red), *slyA*\* (dark green), *slyA*\* expressing ArcZ* (orange), and *slyA*\* expressing wild-type ArcZ (light green). Data are presented as mean ± SD from four independent biological replicates. The red arrow indicates the time point used for reporter quantification in Fig. 2C. (C) AlphaFold3 structural models of the *slyA* mRNA 5′ UTR alone (left) and in complex with ArcZ (right). The *slyA* mRNA 5′ UTR is shown in dark green, the RBS in bright green, and ArcZ in red.

**Figure S3.**
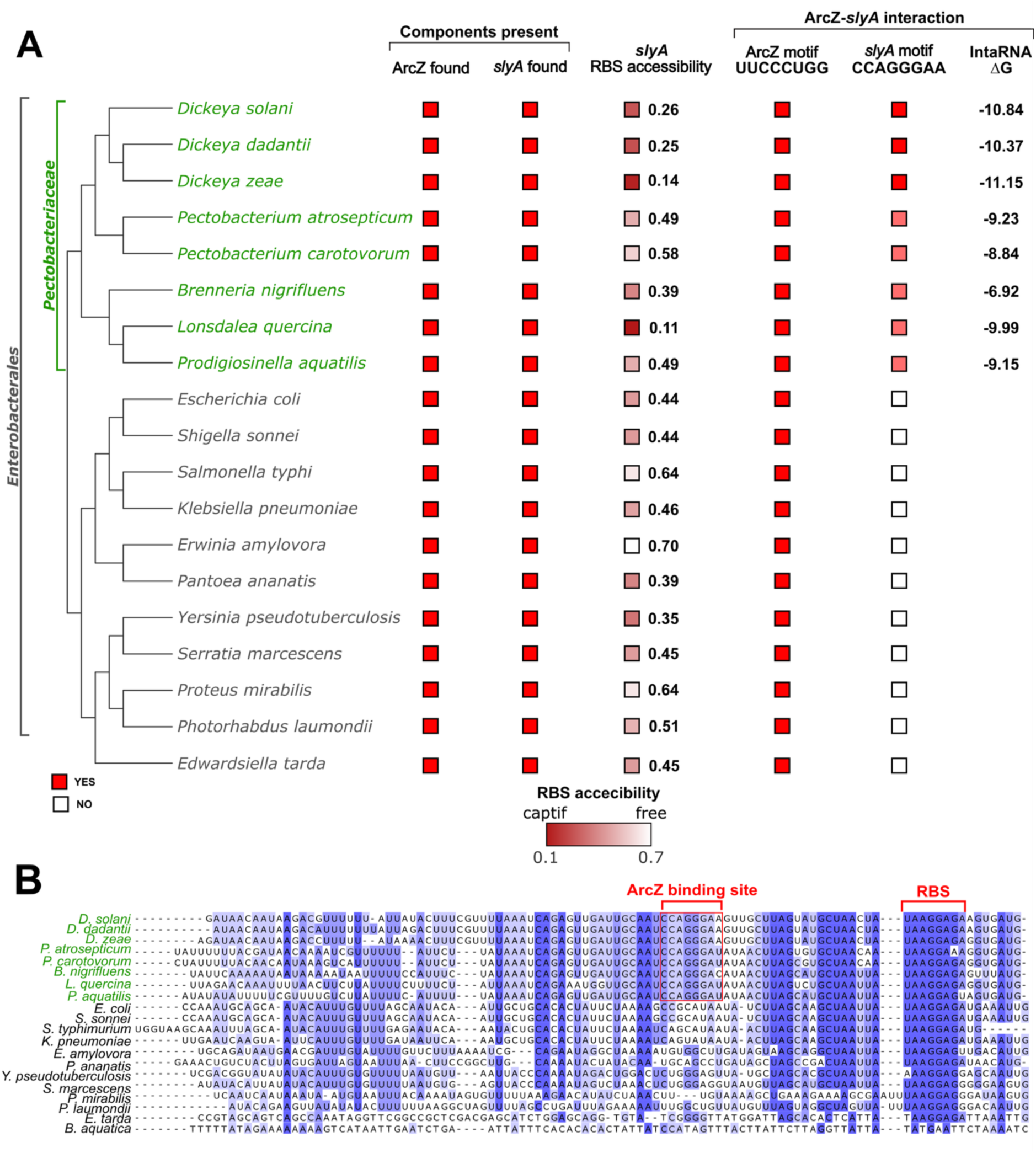
Phylogenetic distribution and sequence conservation of the ArcZ-*slyA* mRNA interaction across *Enterobacterales*. (A) Phylogenetic tree of representative *Enterobacterales* shown together with the distribution of ArcZ, *slyA*, and sequence features associated with the ArcZ-*slyA* mRNA interaction. Species belonging to *Pectobacteriaceae* are indicated in green. Presence or absence of ArcZ and *slyA* is indicated by red and white squares, respectively. Predicted accessibility of the *slyA* RBS in the unbound mRNA is shown according to the indicated color scale, ranging from low accessibility (dark red) to high accessibility (white). Presence of the ArcZ motif (UUCCCUGG) and the complementary *slyA* motif (CCAGGGAA) is indicated by red squares, bright red indicates 100% conservation of the motif, whereas paler red indicates lower conservation. For species in which the complementary *slyA* motif was identified, the predicted ArcZ-*slyA* mRNA interaction energy calculated with IntaRNA is reported as ΔG. (B) Multiple sequence alignment of the *slyA* mRNA 5′ UTR from the indicated *Enterobacterales*. The ArcZ-binding site and ribosome-binding site (RBS) are indicated by red brackets above the alignment. Blue shading indicates nucleotide conservation. Bacterial species in the *Pectobacteriaceae* family are indicated in green.

### Transcriptomic and quantitative proteomic profiling of Δ*slyA* strain reveals concerted control of the T6SS and solanimycin loci

To determine the downstream regulon governed by SlyA and test whether SlyA plays an important role in the antimicrobial pathways affected in Δ*arcZ* cells, we constructed a Δ*slyA* mutant. We compared it to the wild-type strain using both integrated quantitative proteomics and transcriptomics (RNA-seq) (Fig. 3; Fig. S4).

To ensure a direct and robust comparison with our previously established Δ*arcZ* proteome (27), quantitative mass spectrometry of the Δ*slyA* mutant was performed under the exact same physiological condition, namely in M63 minimal medium supplemented with 1% sucrose at early exponential phase (OD_600_ = 0.3) (Fig. 3A) (Table S2). In the Δ*slyA* proteome, proteins associated with both the T6SS and solanimycin pathways were strongly depleted, with the T6SS displaying particularly broad effects across structural, regulatory, and effector components. Several key structural, enzymatic, and regulatory components that were abundant in wild-type cells became completely undetectable in the Δ*slyA* mutant, reaching the technical ratio floor (log_2_FC = -6.64) (Fig. 3A). Within the T6SS, this absolute loss affected both the T6SS sigma factor VasH and the tube protein HcpD, while additional structural and effector elements (TssB, TssC, TssK, TagO, and the spike protein VgrGA) were markedly depleted (Fig. 3A). Concurrently, within the solanimycin pathway, the core non-ribosomal peptide synthetase mega-enzyme SolF was not detected in Δ*slyA* extracts. The co-initiating synthetase SolG (log_2_FC = -3.36) and tailoring enzymes SolC and SolI were also severely depleted (Fig. 3A) (Table S2).

To determine whether the strong proteomic changes observed in Δ*slyA* were accompanied by altered transcription of the corresponding loci, we additionally analyzed the Δ*slyA* transcriptome by RNA-seq. Transcriptomic profiling was performed in LB medium at early exponential phase (OD_600_ = 0.5; Fig. 3B; Fig. S4). Despite the different growth conditions used for proteomic and transcriptomic analyses, the RNA-seq dataset revealed the same broad functional signature. Within the T6SS, transcript depletion encompassed all discrete genomic loci, including the master regulatory activator gene *vasH* (log_2_FC = -2.96), the primary tube gene *hcpB* (log_2_FC = -3.6), the spike gene *vgrGB* (log_2_FC = -3.8), the contractile sheath gene tssB (log_2_FC = -2.9), and multiple *rhsA/C* effector genes (log_2_FC = -2.7 and log_2_FC = -1.4) (Fig. 3B; Fig. S4A). In contrast, within the *sol* cluster only the first gene of the operon, *solA*, was significantly repressed (log_2_FC = -1.5), the remaining sol genes falling below the significance threshold (Fig. 3B; Fig. S4B) (Table S3).

**Figure 3.**
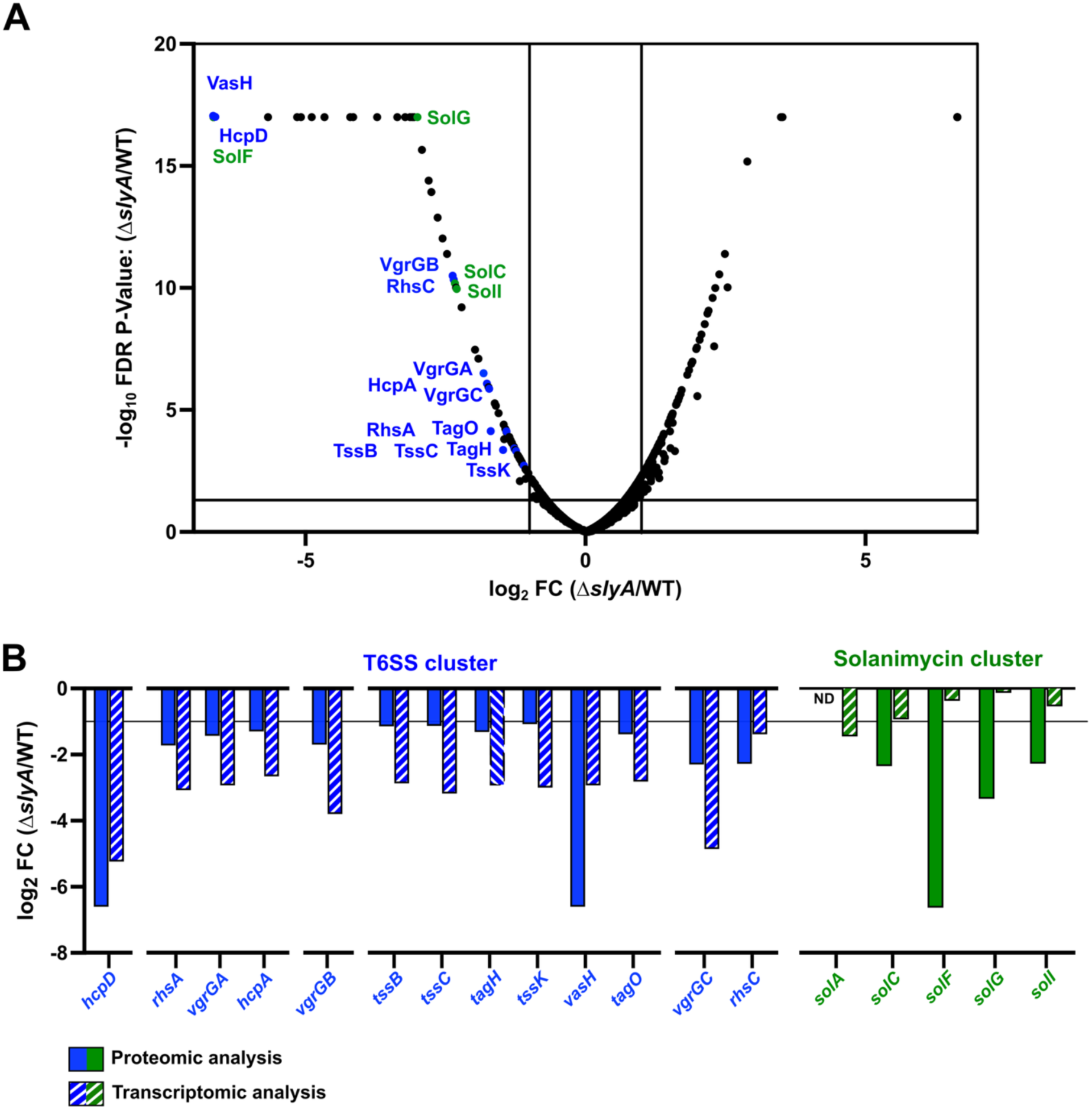
Proteomic and transcriptomic profiling reveals SlyA-dependent control of the T6SS and solanimycin loci. (A) Volcano plot of quantitative proteomic analysis comparing Δ*slyA* and wild-type *D. solani* cells grown in M63 minimal medium supplemented with 1% sucrose at 30°C to OD_600_ = 0.3. The x-axis shows the log_2_ protein abundance ratio (Δ*slyA*/WT), and the y-axis shows −log_10_ P value. Differentially abundant proteins were defined by log_2_ FC ≤ -1 or ≥ 1 and p-value ≤ 0.05. Solanimycin-associated proteins are highlighted in green and T6SS-associated proteins in blue; selected proteins are labeled. Data are from three independent biological replicates. (B) Comparison of proteomic and transcriptomic changes for T6SS- and solanimycin-associated components quantified in the Δ*slyA* mutant. Solid bars represent log_2_ fold changes in protein abundance in Δ*slyA* relative to WT, whereas hatched bars represent the corresponding log_2_ fold changes in transcript abundance determined by RNA-seq. T6SS-associated genes and proteins are shown in blue and solanimycin-associated genes and proteins in green. RNA-seq was performed on cells grown in LB medium at 30°C to OD_600_ = 0.5. The horizontal line indicates a log_2_ fold change of -1. ND indicates components not detected in the corresponding dataset. Proteomic and transcriptomic data are each derived from three independent biological replicates.

**Figure S4.**
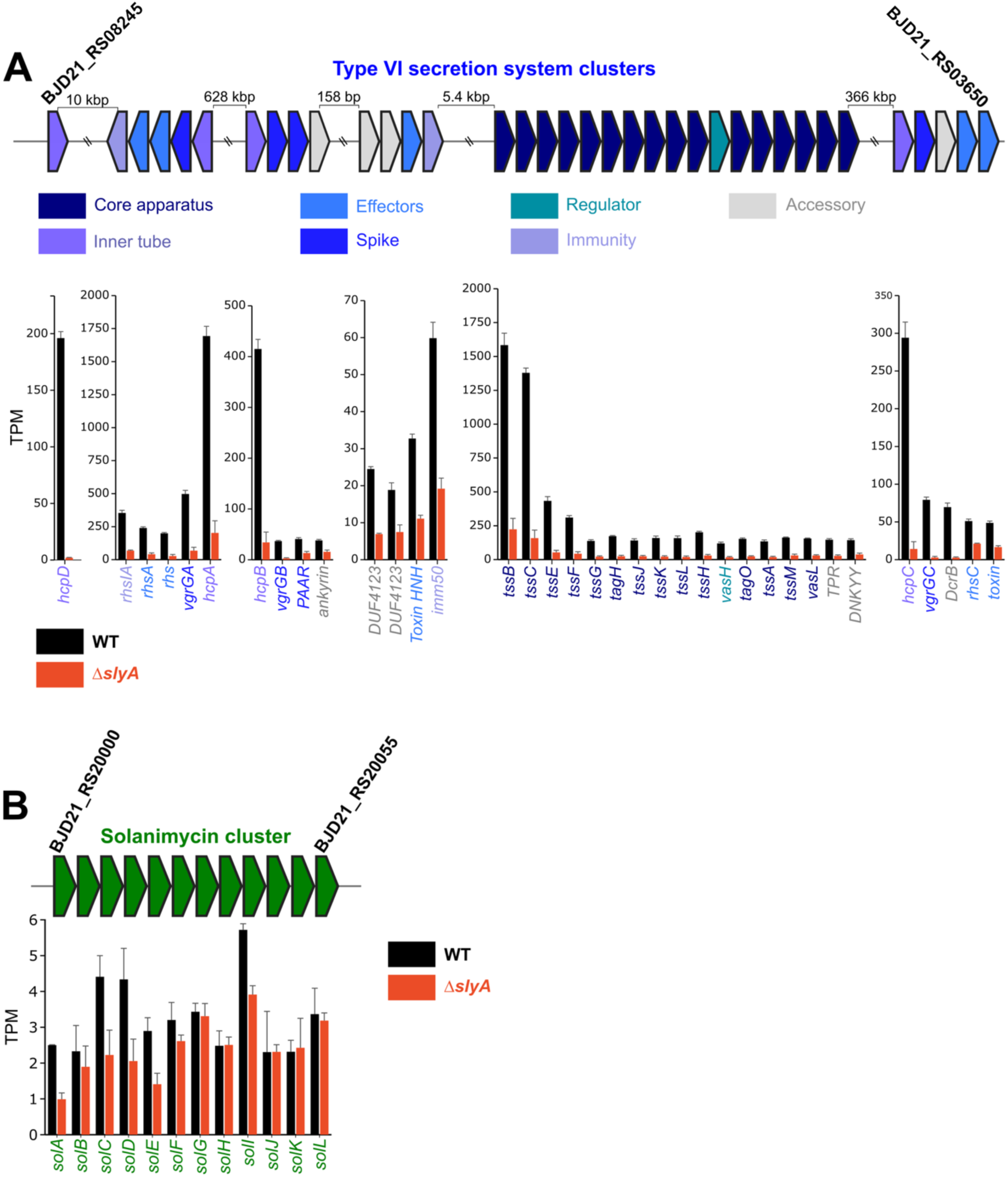
Expression profiles across the complete T6SS and solanimycin loci in wild-type and Δ*slyA* cells. (A) Genomic organization and RNA-seq expression profiles of the *D. solani* T6SS-associated loci. Genes are represented by arrows indicating transcriptional orientation and are color-coded according to functional category: core apparatus (dark blue), inner tube (purple), spike (blue), effectors (light blue), immunity proteins (light purple), regulators (teal), and accessory proteins (grey). Bar graphs show transcript abundance expressed as transcripts per million (TPM) for wild-type (WT, black) and Δ*slyA* (red) cells. (B) Genomic organization and RNA-seq expression profiles of the solanimycin biosynthetic cluster. *sol* genes are shown in green. Bar graphs show TPM values for WT (black) and Δ*slyA* (red) cells. Cells were grown in LB medium at 30°C to OD_600_ = 0.5. Data are presented as mean ± SD from three independent biological replicates.

Together, the proteomic and transcriptomic datasets identify SlyA as a major positive regulator of both antimicrobial systems, with loss of SlyA causing broad depletion of T6SS components and reduced expression of the solanimycin biosynthetic pathway. The concordant changes at the transcript and protein levels further suggest that SlyA acts predominantly through transcriptional control of these loci.

### SlyA directly binds and activates regulated solanimycin and T6SS promoters

To determine whether SlyA directly controls the antimicrobial loci identified by the omics analyses, we combined transcriptional promoter-*lux* fusions with electrophoretic mobility shift assays (EMSAs) using purified SlyA.

Transcriptional *luxCDABE* reporter fusions demonstrated strong SlyA-dependent activity for three representative promoters: P*_solA_*, P*_hcpA_*, and P*_hcpB_* (Fig. 4A). Bioluminescence driven by each of these promoters was reduced in both the Δ*arcZ* and Δ*slyA* backgrounds, and expression of *slyA in trans* restored promoter activity in Δ*slyA* cells (Fig. 4A). Hcp proteins constitute the inner tube of the T6SS injection apparatus and are required for effector delivery, making *hcpA* and *hcpB* informative indicators of T6SS expression. In contrast, while *tssB* and *vgrGB* transcripts were reduced in the Δ*slyA* RNA-seq dataset (Fig. 3B; Fig. S4A), the corresponding P*_tssB_* and P*_vgrGB_* promoter fusions retained wild-type activity in Δ*slyA* cells, although both were reduced in the Δ*arcZ* strain (Fig. S5A). Thus, the inability of isolated P*_tssB_* and P*_vgrGB_* fragments to recapitulate native SlyA dependence suggests that regulation at these loci involves indirect mechanisms, distal cis-acting elements or chromosomal context not captured by the reporter constructs.

We next evaluated whether SlyA binds directly to the promoters that exhibited SlyA-dependent activation *in vivo*. In EMSAs, purified His_6_-SlyA (Fig. S5B) caused concentration-dependent electrophoretic mobility shifts of the P*_solA_*, P*_hcpA_*, and P*_hcpB_* DNA fragments, whereas no binding was observed with a non-specific control fragment (Fig. 4B). Quantification of the bound fraction revealed comparable apparent half-maximal binding concentrations for the three target promoters (Fig. 4C).

Together with the *in vivo* reporter data, these findings demonstrate that SlyA directly binds and activates selected promoters driving solanimycin biosynthesis and T6SS tube assembly, confirming a direct mechanism of transcriptional control for both antimicrobial systems.

**Figure 4.**
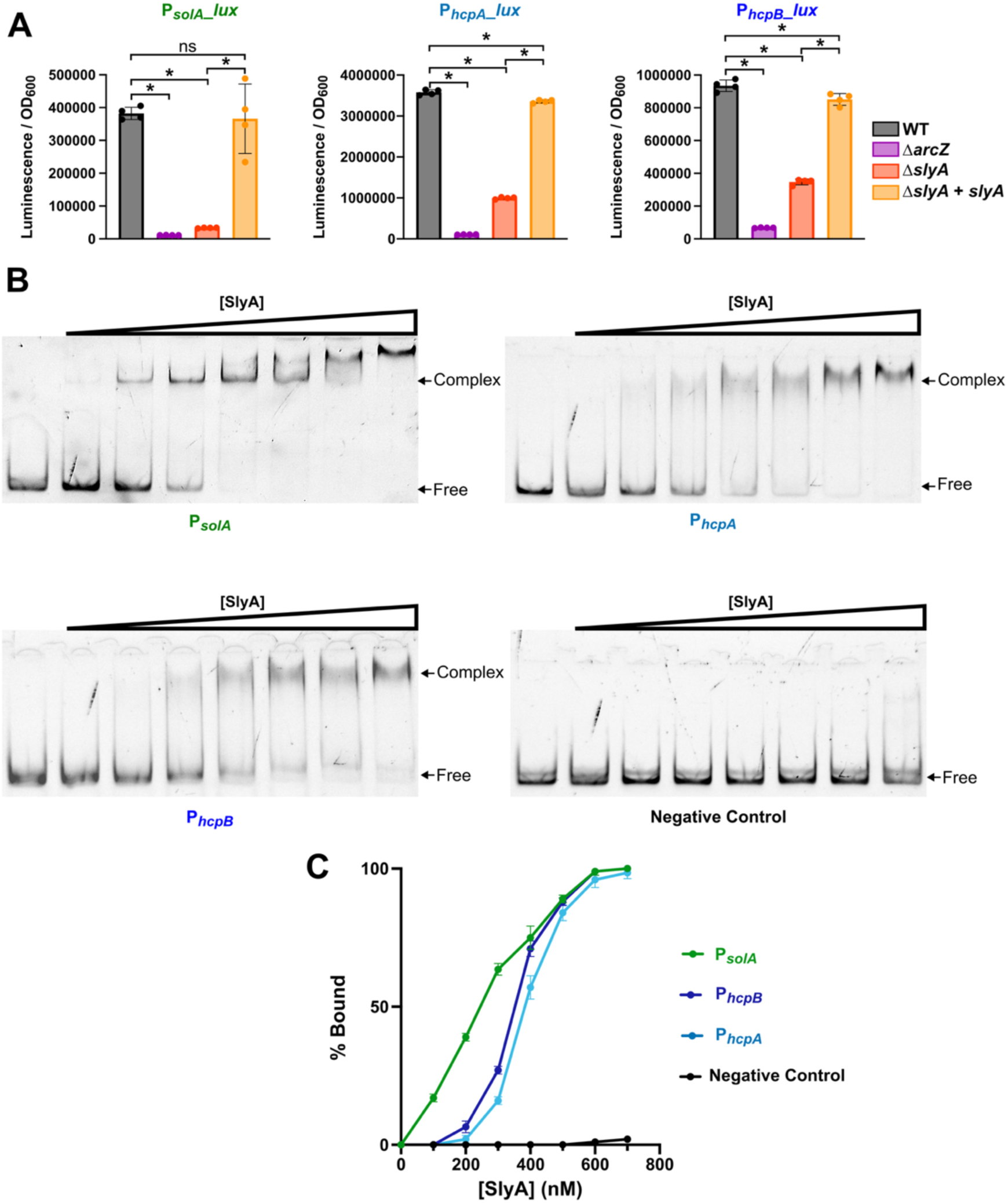
SlyA directly binds and activates selected solanimycin and T6SS promoters. (A) Activity of transcriptional lux fusions to the *solA*, *hcpA*, and *hcpB* promoters (P*_solA_*, P*_hcpA_*, and P*_hcpB_*) in the indicated genetic backgrounds (WT, Δ*arcZ*, Δ*slyA*, Δ*slyA* complemented *slyA*^+^). Promoter activity was monitored at OD_600_ = 0.3 as luminescence normalized to OD_600_. Bars represent mean ± SD from four independent biological replicates; individual replicates are shown as dots. Statistical comparisons were performed using pairwise Mann-Whitney tests. *P < 0.05; **P < 0.01; ***P < 0.001; ns, not significant. (B) Electrophoretic mobility shift assays (EMSAs) of purified His_6_-SlyA with 10 nM DNA fragments of P*_solA_* (150 bp), P*_hcpA_* (158 bp), P*_hcpB_* (171 bp). DNA fragments were incubated with the indicated concentrations of purified SlyA (0, 100, 200, 300, 400, 500, 600 and 700 nM monomer SlyA). A control DNA fragment was included as a negative binding control (162 bp). Free and SlyA-bound DNA species are indicated. P*_solA_*, P*_hcpA_*, and P*_hcpB_* and negative control are shown in green, light blue, dark blue, and black respectively. (C) Quantification of SlyA binding to P*_solA_*, P*_hcpA_*, P*_hcpB_* and negative control from the EMSAs shown in panel B. The fraction of bound DNA is plotted as a function of SlyA concentration (nM). Curves represent fits to the experimental data. P*_solA_*, P*_hcpA_*, P*_hcpB_* and negative control are shown in green, light blue, dark blue, and dark respectively. Data are presented as mean ± SD from two independent experiments.

**Figure S5.**
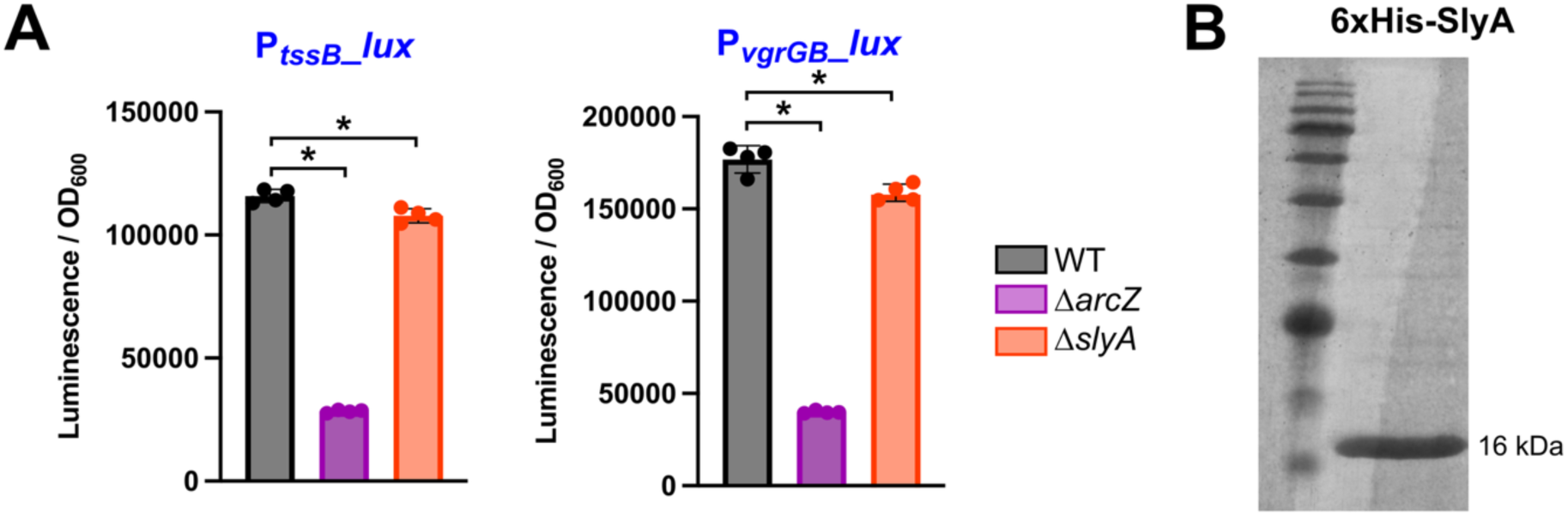
Additional T6SS promoter activities and purification of His_6_-SlyA. (A) Activity of transcriptional *lux*CDABE fusions to the *tssB* and *vgrGB* promoters (P*_tssB_* and P*_vgrGB_*) in wild-type (WT), Δ*arcZ*, and Δ*slyA D. solani* cells. Promoter activity was monitored at OD_600_ = 0.3 as luminescence normalized to OD_600_. WT, Δ*arcZ*, and Δ*slyA* strains are shown in grey, purple, and orange, respectively. Bars represent mean ± SD from four independent biological replicates; individual replicates are shown as dots. Statistical comparisons were performed using pairwise Mann-Whitney tests. *P < 0.05; **P < 0.01; ***P < 0.001; ns, not significant. (B) SDS-PAGE analysis of purified His_6_-SlyA used for electrophoretic mobility shift assays. The position of His_6_-SlyA at approximately 16 kDa is indicated.

### The ArcZ-SlyA cascade controls T6SS-mediated killing and solanimycin-dependent antifungal activity

To determine whether the ArcZ-SlyA regulatory cascade governs the functional antimicrobial activities encoded by the T6SS and solanimycin pathways, we evaluated interbacterial and antifungal antagonism in wild-type and mutant strains.

In interbacterial competition assays against the potato pathogen *P. atrosepticum*, wild-type *D. solani* reduced prey survival by approximately 1,000-fold relative to the T6SS-deficient Δ*tssM* mutant (Fig. 5A; Fig. S6C). Deletion of either *arcZ* or *slyA* similarly impaired interbacterial killing to levels comparable to those observed in the Δ*tssM* strain, and killing was restored upon genetic complementation with the corresponding wild-type gene (Fig. 5A; Fig. S6C; Fig. S7B and C). These differences were not attributable to altered initial competitor-to-prey ratios or growth defects under the conditions tested (Fig. S6A; Fig. S7A). Thus, these assays demonstrate that both ArcZ and SlyA are required for T6SS-dependent killing of *P. atrosepticum*.

In parallel, antifungal activity was evaluated against the ascomycete yeast *K. lactis*. Wild-type *D. solani* produced a measurable growth inhibition zone, whereas antifungal activity was abolished in Δ*arcZ*, Δ*slyA*, and solanimycin-deficient Δ*sol* strains (Fig. 5A; Fig. S6E). Antifungal activity was restored upon complementation of Δ*arcZ* and Δ*slyA* mutants with the corresponding wild-type gene in *trans* (Fig. 5A; Fig. S6E). The phenotypic correspondence among Δ*arcZ*, Δ*slyA*, and Δ*sol* strains identifies solanimycin production as the primary driver of this antifungal phenotype. Together, these assays confirm that the ArcZ-SlyA cascade controls two mechanistically distinct antimicrobial outputs: T6SS-dependent antibacterial killing and solanimycin-dependent antifungal activity.

To test whether these phenotypes depend specifically on direct ArcZ-*slyA* base pairing, we evaluated the compensatory-mutation system. Initial competitor-to-prey ratios were comparable across all strains of this set (Fig. S6B). Disruption of the ArcZ target site in the *slyA* mRNA 5′ UTR (*slyA*\*) impaired both T6SS-mediated killing and antifungal activity, and expression of wild-type ArcZ failed to restore either output (Fig. 5B; Fig. S6D and F). In contrast, co-expression of the compensatory ArcZ* variant, which restores complementarity with *slyA*\*, rescued both interbacterial killing and antifungal inhibition (Fig. 5B; Fig. S6D and F).

These allele-specific genetic rescues establish direct ArcZ-*slyA* mRNA base pairing as the causal regulatory link through which ArcZ, via SlyA, controls both T6SS-dependent antibacterial killing and solanimycin-dependent antifungal activity in *D. solani*.

**Figure 5.**
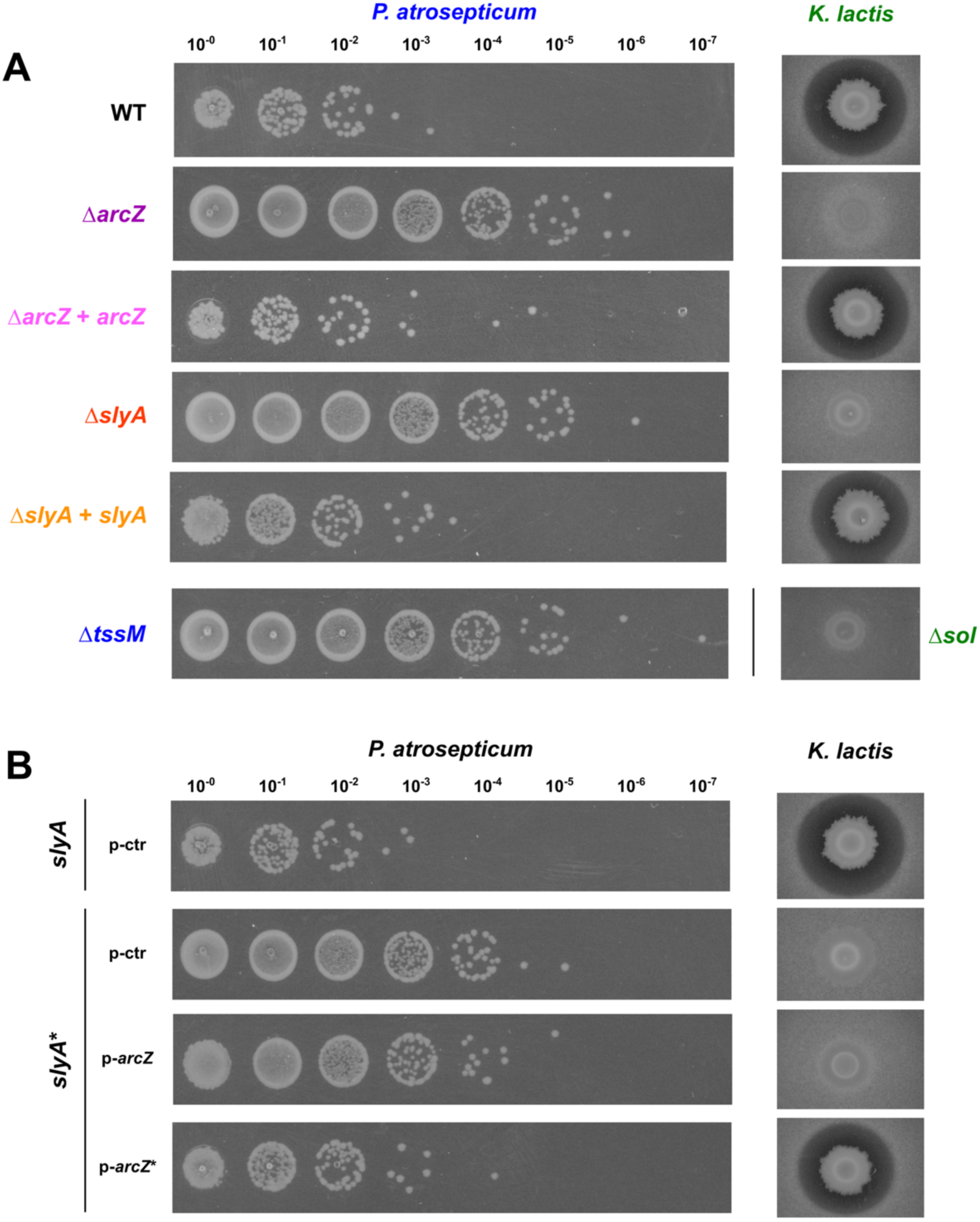
The ArcZ-SlyA cascade controls T6SS-mediated antibacterial killing and solanimycin-dependent antifungal activity. (A) Antibacterial and antifungal activities of the indicated *D. solani* strains: (WT), *ΔarcZ*, Δ*arcZ* complemented with plasmid-borne *arcZ*, Δ*slyA*, Δ*slyA* complemented with plasmid-borne *slyA*, and a negative control strain (the T6SS-deficient Δ*tssM* strain on the left, and the solanimycin-deficient Δ*sol* strain on the right). Left: serial-dilution plating of the potato pathogen *P. atrosepticum* recovered after 16-hour interbacterial competition assays against the indicated *D. solani* strains. Right: inhibition assays of *K. lactis* by the indicated *D. solani* strains. Representative images are shown; quantitative analyses are provided in Fig. S6C and E. (B) Antibacterial and antifungal activities of *D. solani* strains carrying the compensatory ArcZ-*slyA* mutation system. The *slyA*\* allele contains substitutions within the ArcZ-binding site, and *arcZ*\* carries the complementary substitutions restoring base-pairing potential. Left: serial-dilution plating of the potato pathogen *P. atrosepticum* recovered after 16-hour interbacterial competition assays against the indicated *D. solani* strains. Right: inhibition assays of *K. lactis* by the indicated *D. solani* strains. Representative images are shown; corresponding quantifications are provided in Fig. S6D and F.

**Figure S6.**
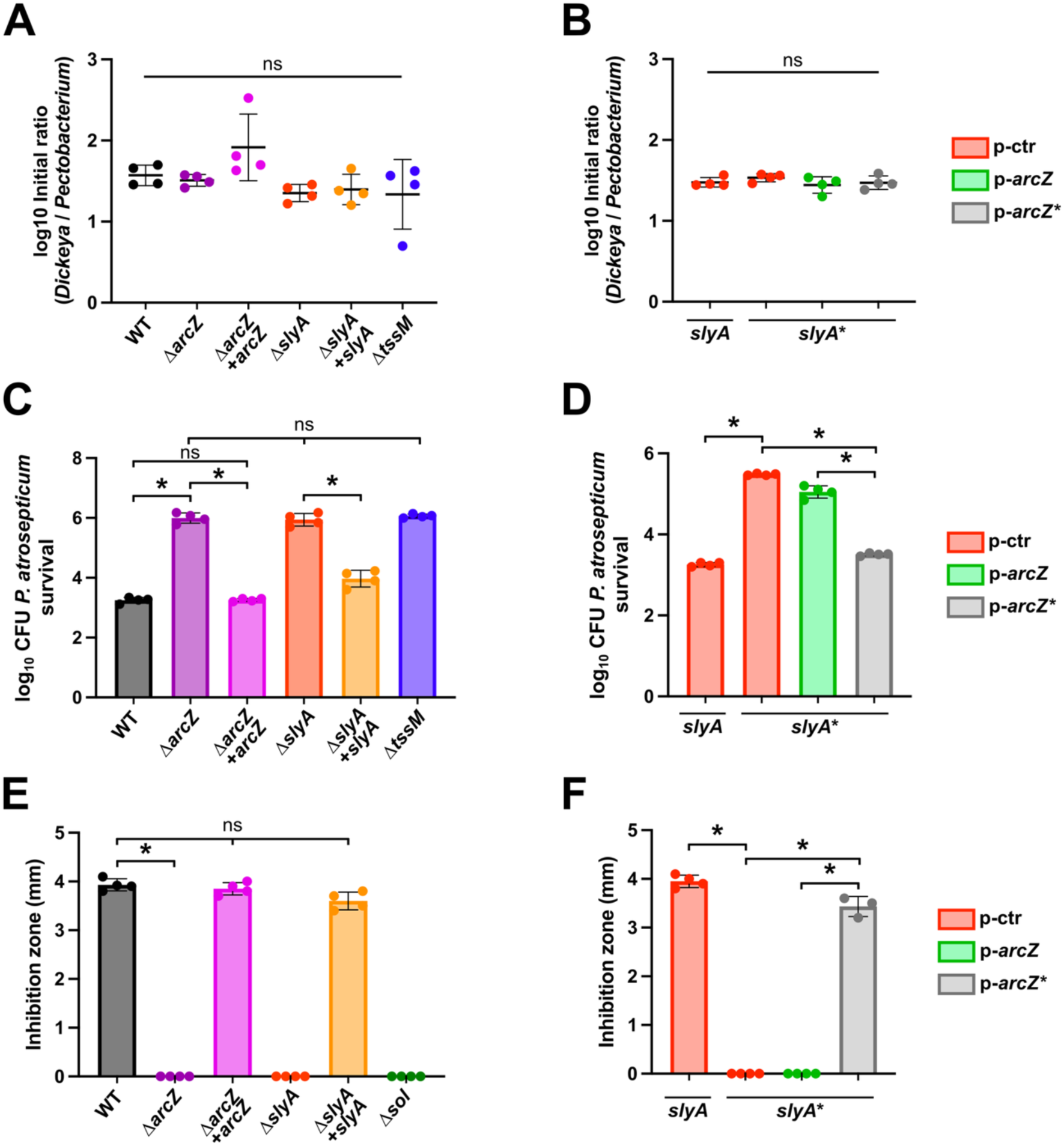
Quantification of ArcZ- and SlyA-dependent antibacterial and antifungal activities. (A, C) Initial competitor-to-prey ratios (A) and prey survival (C) in interbacterial competition assays between the indicated *D. solani* strains (WT, Δ*arcZ*, Δ*arcZ* complemented with plasmid-borne *arcZ*, Δ*slyA*, Δ*slyA* complemented with plasmid-borne *slyA*, and Δ*tssM*) and *P. atrosepticum*. Ratios are expressed as log_10_ *D. solani* / *P. atrosepticum* for panel A. (B, D) Initial competitor-to-prey ratios (B) and prey survival (D) in interbacterial competition assays between *D. solani* strains carrying the ArcZ-*slyA* compensatory-mutation system and *P. atrosepticum*. *D. solani* strains carry the wild-type *slyA* or *slyA*\* allele and contained either an empty control plasmid (p-ctr, red), a plasmid bearing wild-type *arcZ* (p-arcZ, green), or a plasmid bearing the compensatory *arcZ*\* allele (p-*arcZ*\*, grey), as indicated. (E, F) Quantification of antifungal activity against *K. lactis* for the indicated *D. solani* strains. Inhibition zones were measured in millimeters. Results for *D. solani* strains WT, Δ*arcZ*, Δ*arcZ* complemented with plasmid-borne *arcZ*, Δ*slyA*, Δ*slyA* complemented with plasmid-borne *slyA*, and Δ*sol* are presented in E. Results for *D. solani* strains carrying the wild-type *slyA* or *slyA*\* allele and containing either an empty control plasmid (p-ctr, red), a plasmid bearing wild-type *arcZ* (p-arcZ, green), or a plasmid bearing the compensatory *arcZ*\* allele (p-*arcZ*\*, grey) are presented in F. Data are presented as mean ± SD from four independent biological replicates; individual replicates are shown as dots. Statistical comparisons were performed using pairwise Mann-Whitney tests. *P < 0.05; ns, not significant.

**Figure S7.**
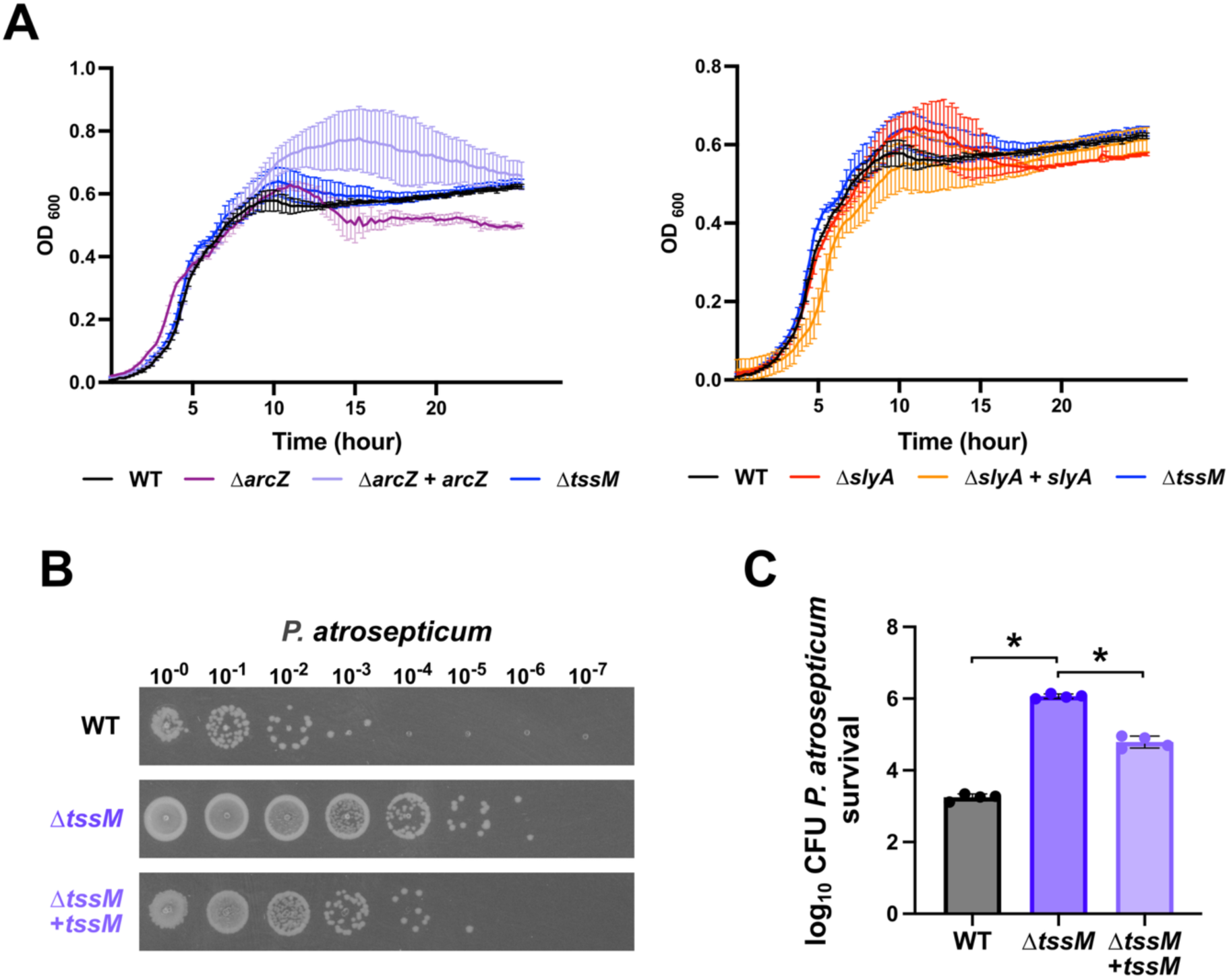
Growth profiles of strains used in competition assays and complementation of the T6SS-deficient Δ*tssM* mutant. (A) Growth curves of *D. solani* wild-type (WT), Δ*arcZ*, Δ*arcZ* complemented with plasmid-borne *arcZ* and Δ*tssM* (left) (in black, purple, light purple and blue respectively) and WT, Δ*slyA*, Δ*slyA* complemented with plasmid-borne *slyA*, and Δ*tssM* strains (right) (in black, red, orange, and blue, respectively). Growth was monitored by OD_600_ over 24 h. Data are presented as mean ± SD from 4 independent biological replicates. (B) Representative serial-dilution plating of *P. atrosepticum* recovered after interbacterial competition with *D. solani* strains WT, Δ*tssM*, or Δ*tssM* complemented with plasmid-borne *tssM*. Dilution factors are indicated above the plates. (C) Quantification of *P. atrosepticum* survival corresponding to the competition assays shown in panel B, expressed as log_10_ CFU. *D. solani* strains WT, Δ*tssM*, and Δ*tssM* complemented with plasmid-borne *tssM* are shown in grey, dark purple, and light purple, respectively. Bars represent mean ± SD; individual biological replicates are shown as dots. Statistical comparisons were performed using pairwise Mann-Whitney tests. *P < 0.05.

## Discussion

In this study, we demonstrate that the Hfq-dependent sRNA ArcZ directly activates the translation of the MarR-family transcription factor SlyA, establishing a two-tier regulatory cascade that coordinates solanimycin-dependent antifungal activity and T6SS-mediated interbacterial killing in *D. solani* (Fig. 6). Our RIL-seq data identify ArcZ as the most highly connected Hfq-dependent sRNA under the tested conditions. Rather than establishing direct base-pairing interactions with each downstream target transcript, ArcZ expands its regulatory breadth by targeting higher-order transcriptional regulators. We show here that, through direct activation of *slyA* translation, ArcZ propagates its post-transcriptional signal across extensive downstream gene networks. This hierarchical architecture provides a mechanistic explanation for the pleiotropic phenotypes associated with ArcZ (11) and a direct mechanistic resolution for how ArcZ activates extensive, unlinked antimicrobial systems (the 40-kb sol cluster and multi-operon T6SS loci) despite the absence of direct *in vivo* ArcZ interactions with their core structural transcripts in the RIL-seq dataset.

**Figure 6.**
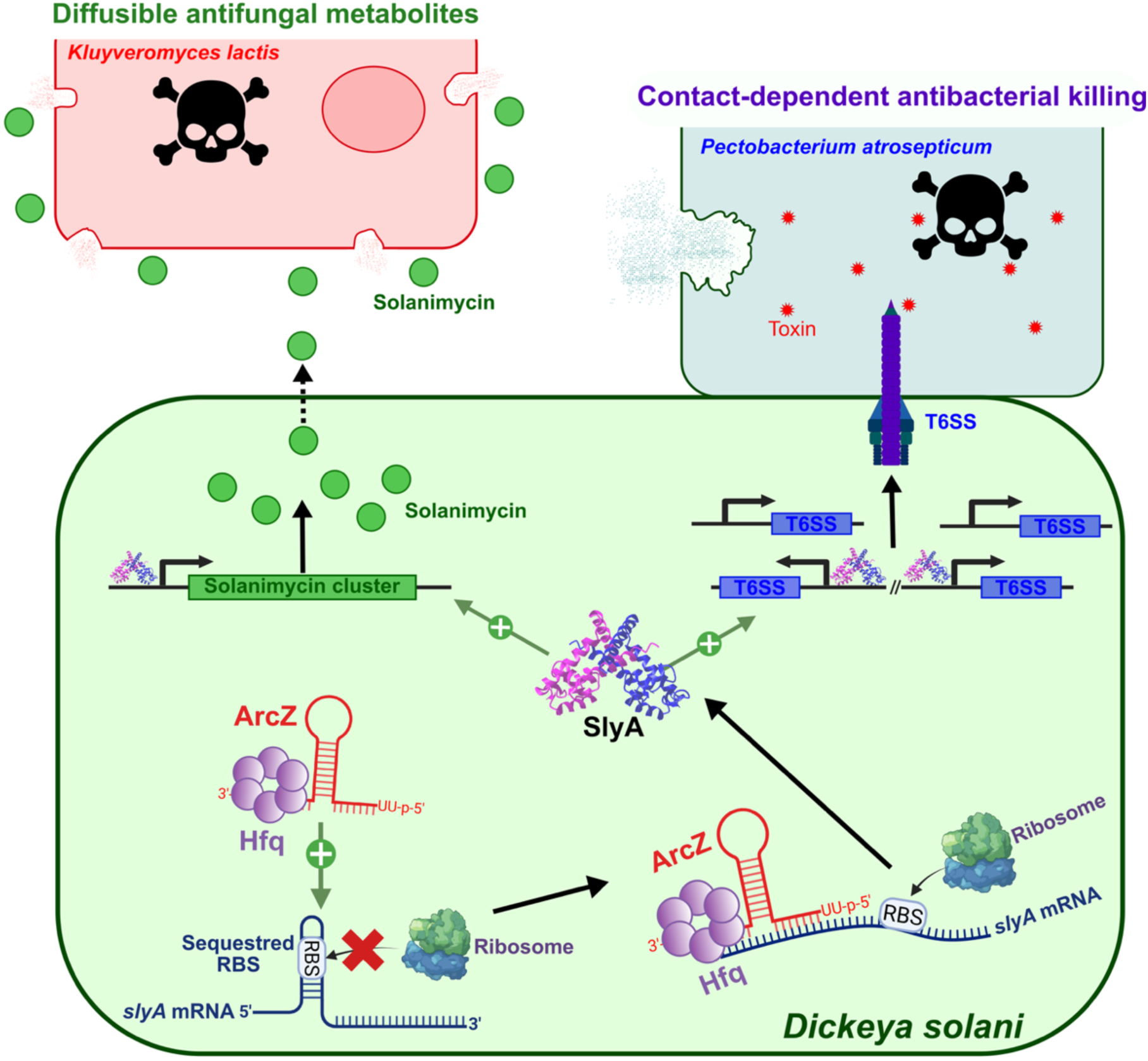
Model for ArcZ-SlyA-dependent coordination of antibacterial and antifungal activities in *D. solani*. Schematic representation of the regulatory cascade identified in this study. ArcZ base-pairs with the 5′ untranslated region of *slyA* mRNA, increasing RBS accessibility and thereby promoting *slyA* mRNA translation. SlyA subsequently activates the transcription of multiple target genes by binding to their promoter regions, including genes of the solanimycin biosynthetic cluster and genes encoding the T6SS.

Solanimycin mediates antifungal activity, illustrated here against the yeast *K. lactis* (red cell), whereas the T6SS mediates antibacterial activity, illustrated here against *P. atrosepticum* (blue cell).

Mechanistically, ArcZ activates *slyA* translation by base-pairing with its 5′ UTR, resolving an inhibitory secondary structure that sequesters the ribosome-binding site (Fig. 2, Fig. 6). This mode of translational activation resembles ArcZ-dependent control of *rpoS*, in which sRNA annealing promotes accessibility of the translation initiation region (29, 30). SlyA represents one branch of a broader regulatory hub controlled by ArcZ. Indeed, the RIL-seq dataset also revealed that *rpoS* mRNA was captured as an ArcZ target at high cell density, and *pecT* mRNA, encoding a major transcriptional repressor of *Dickeya* virulence, was among the most abundant ArcZ interaction partners. Direct repression of *pecT* by ArcZ was previously characterized in *D. dadantii* (31). Consistent with these regulatory links, decreased RpoS abundance and elevated PecT levels were initially detected in the *ΔarcZ* proteomic analysis from our previous study (27). Thus, ArcZ appears positioned to coordinate several transcriptional regulators controlling distinct components of the *D. solani* pathogenic and competitive program, with SlyA providing the direct regulatory connection to the antimicrobial systems characterized here. Furthermore, the marked depletion of structural and enzymatic components required for antifungal and antibacterial activities in the Δ*slyA* proteome, including VasH, HcpD, and SolF, supports a central role for SlyA in propagating the upstream ArcZ signal to antimicrobial systems. This predominantly hierarchical mode of regulation does not, however, exclude direct control of individual downstream components. The T6SS-associated *vgrGB* transcript was also recovered as a direct ArcZ interaction partner in the RIL-seq dataset, suggesting that ArcZ may combine broad regulation through SlyA with direct post-transcriptional regulation of selected T6SS transcripts (Table S1).

In *Pectobacteriaceae*, *slyA* transcription is integrated into a multilayered network that responds to host and environmental stress. Transcription of *slyA* is stimulated by the oxidative stress regulator OhrR (20) and by the PhoP-PhoQ two-component system (23), while being directly repressed by the redox-sensing ArcAB two-component system (21). SlyA activity is also connected to quorum sensing and cyclic-di-GMP signaling through the AraC-family regulator VfmE (32). Our results add a post-transcriptional layer to this network by showing that SlyA production is directly controlled by ArcZ. The ArcZ sequence involved in this interaction is broadly conserved, whereas the complementary sequence in the *slyA* 5′ UTR is restricted to *Pectobacteriaceae* (Fig. 2E; Fig. S3). Predicted accessibility of the *slyA* ribosome-binding site varies among members of this family, with lower accessibility in the *Dickeya* sequences examined here than in representative *Pectobacterium* sequences. Although these predictions require experimental validation, they raise the possibility that the quantitative contribution of ArcZ to SlyA production differs among *Pectobacteriaceae*. Testing whether ArcZ post-transcriptionally controls SlyA accumulation and downstream competitive determinants in representative *Pectobacterium* strains will be an informative direction for future studies to determine the functional conservation of this checkpoint across the *Pectobacteriaceae*.

Our data identify both the solanimycin cluster and the T6SS as direct components of the SlyA regulon. SlyA binds the P*_solA_*, P*_hcpA_* and P*<u>_hcpB_</u>* promoter regions *in vitro*, and transcription from these promoters is reduced in the absence of SlyA (Fig. 4). This direct promoter binding is consistent with the broader role of SlyA-family regulators in controlling horizontally acquired and H-NS-silenced loci. SlyA proteins recognize relatively degenerate AT-rich sequences and can activate transcription either through conventional promoter regulation or by counteracting H-NS-mediated silencing (14, 15, 33). Such a mechanism is plausible for antimicrobial loci, which are frequently associated with horizontally acquired genomic regions. However, although our data demonstrate direct promoter binding and activation by SlyA, establishing whether this activation involves relief of H-NS-mediated silencing will require *hns* epistasis experiments and comparative mapping of SlyA and H-NS occupancy *in vivo*.

The regulation of the T6SS by SlyA also integrates into a regulatory framework previously described in *Pectobacteriaceae*. In *P. atrosepticum*, transcription of the T6SS core cluster and auxiliary *hcp*-*vgrG* operons requires the alternative sigma factor σ^54^ (RpoN) and its cognate enhancer-binding protein VasH (34). This architecture appears to be conserved in *D. solani*: the canonical σ^54^ −24/−12 promoter elements are found upstream of the corresponding hcp modules, including P*_hcpA_* and P*_hcpB_*, at which we demonstrate direct SlyA binding (Fig. S8A), and the structural T6SS cluster is colinear with that of *P. atrosepticum* and encodes a *vasH* orthologue at the same position (Fig. S8B). VasH abundance is strongly reduced in the Δ*slyA* proteome (Fig. 3A). SlyA may control T6SS expression at two levels, through direct regulation of hcp promoters and through its effect on *vasH* expression. Whether SlyA and Eσ^54^-VasH cooperate at these promoters remains to be determined.

SlyA DNA binding is itself antagonized by aromatic carboxylates (35, 36). Among these, precursors of catecholate siderophores and of quinone electron carriers inhibit SlyA, linking its activity to iron availability and respiratory status (36). SlyA is therefore positioned to integrate metabolic information into the transcriptional control of antimicrobial systems.

Previous studies indicate that T6SS expression in *Dickeya* and *Pectobacterium* is strongly influenced by environmental and host-associated conditions. In *P. atrosepticum* SCRI1043, potato extracts stimulate expression of multiple *hcp* genes and *vgrG* and induce components of the T6SS, while T6SS-associated genes are also subject to quorum-sensing regulation (37–39). In *D. dadantii*, T6SS genes are differentially expressed during plant colonization and are controlled by the global regulator PecS, with the T6SS biosynthetic genes, multiple *hcp* genes and *vgrG* being derepressed in a pecS mutant (40). Similarly, the complete *tss* cluster and several *hcp*-*vgrG*-associated islands of *P. brasiliense* PBR1692 are strongly induced during potato tuber infection (41). These observations establish that T6SS deployment in *Pectobacteriaceae* is integrated into regulatory programs responding to host-associated conditions, cell density and infection stage. Our identification of SlyA and ArcZ as upstream regulators in *D. solani* adds a post-transcriptional component to this regulatory landscape. SlyA is also required for antibacterial activity in *P. brasiliense*, although through a different weapon: deletion of *slyA*, of fur or of *expI* abolishes carbapenem production, itself dependent on environmental iron and oxygen, and putative Fur boxes are predicted upstream of *slyA* in several *Pectobacterium* genomes (42). Across *Pectobacteriaceae*, SlyA therefore appears to control whichever antibacterial biosynthetic locus a given genome carries, carbapenem in *P. brasiliense*, solanimycin and the T6SS in *D. solani*, a pattern consistent with the metabolic responsiveness of SlyA described above.

Despite the widespread occurrence and regulated expression of T6SS loci in *Pectobacteriaceae*, direct demonstrations of their contribution to interbacterial killing remain scarce. In *D. dadantii* 3937, the RhsA and RhsB proteins mediate contact-dependent inhibition of neighboring cells, and RhsB-dependent competition requires the associated VgrG proteins, supporting delivery through a T6SS-related mechanism (43). In *P. brasiliense* PBR1692, a T6SS-deficient mutant loses the ability to reduce the survival of *D. dadantii*, *D. chrysanthemi* and *P. carotovorum* by 2 to 3 log in potato tubers, a defect restored by complementation, whereas the same mutant competes as efficiently as the wild type in co-culture, where killing depends instead on carbapenem production (42). The antibacterial contribution of the *P. brasiliense* T6SS is therefore conditional on the plant environment and on the identity of the competitor (42).

We show here that the *D. solani* T6SS mediates interbacterial killing under standard laboratory conditions, in the absence of exogenous plant-derived signals. Unlike the *P. brasiliense* system (42), the *D. solani* T6SS therefore displays antibacterial activity outside the plant environment. Whether plant-derived signals further enhance T6SS activity in *D. solani* remains to be determined.

**Figure S8.**
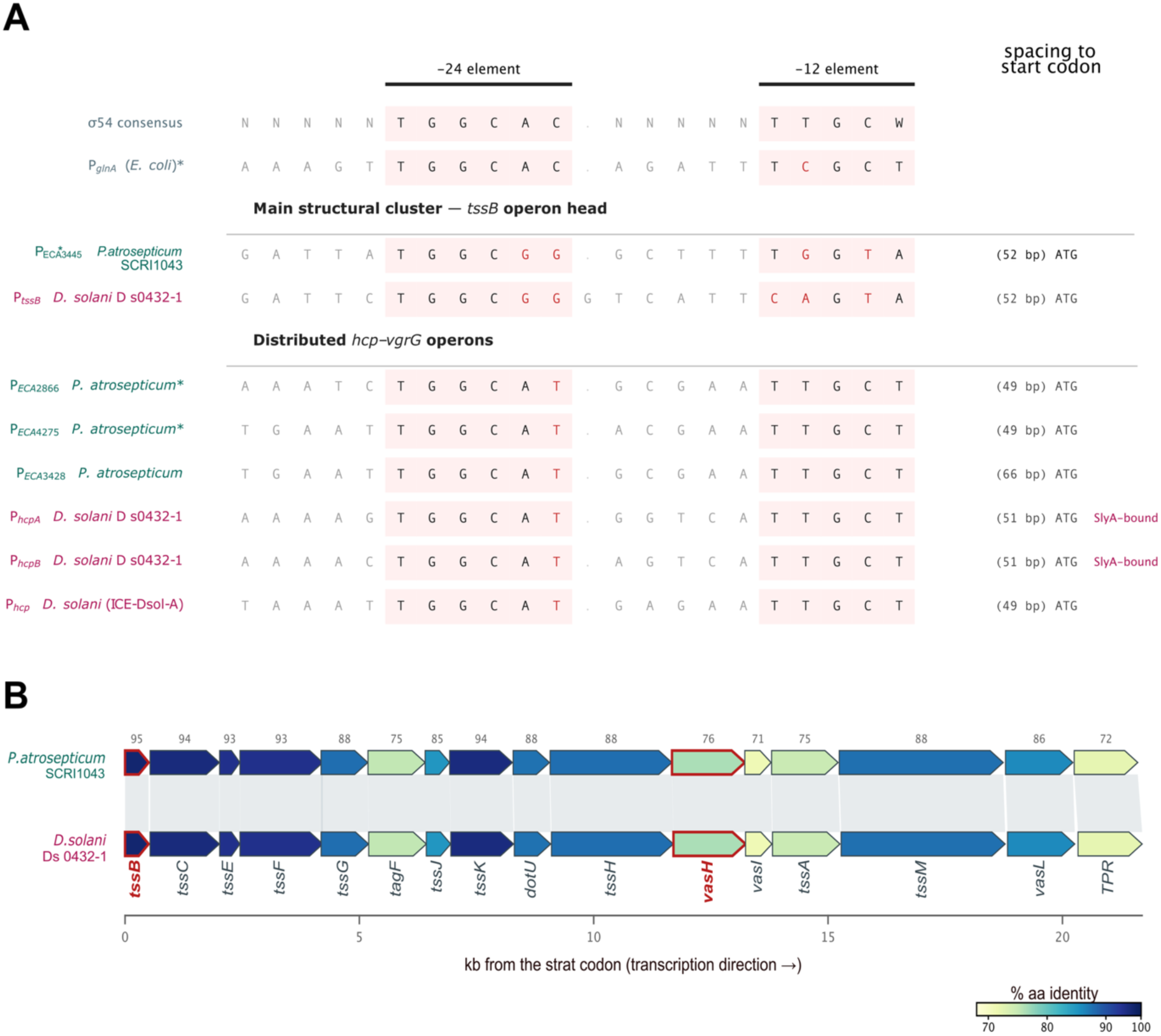
Conservation of the putative σ^54^ promoter elements and of the main T6SS structural cluster between *P. atrosepticum* SCRI1043 and *D. solani* D s0432-1. (A) Promoter regions aligned on the -24 and -12 elements, following the presentation of Fig. 1 of ref (34). Each row shows 5 nt of upstream context, the -24 element, the variable spacer (grey dots denote alignment gaps), the -12 element, and the number of base pairs separating the 3’ end of the box from the start codon. Bases departing from the σ^54^ consensus (TGGCAC-N5-TTGCW) are in red. Block A, head of the main structural cluster; block B, distributed *hcp-vgrG* operons. Asterisks mark promoters shown to be retarded by σ^54^ and VasH *in vitro* in ref. (34). All spacings were computed from the current assemblies (BX950851.1, NZ_CP017453.1). (B) Synteny of the main structural cluster. Arrows are genes drawn to scale in the direction of transcription, aligned on the *tssB* start codon; the sixteen pairs shown are reciprocal best BLASTP hits and are shaded by amino-acid identity (71-95%, median 88%), with connectors joining orthologues. The cluster spans 21,606 bp in *P. atrosepticum* and 21,699 bp in *D. solani* with no insertion or rearrangement, and the enhancer-binding protein VasH (red outline) occupies the same position in both.

This killing activity can be placed alongside epidemiological observations from potato. Interactions between *Dickeya* and *Pectobacterium* species are ecologically plausible during potato infection, as field surveys have documented frequent co-occurrence of *D. dianthicola* and *P. parmentieri* within naturally infected potato material, with both species detected in 20.6% of samples (44). In a fourteen-year survey of Finnish seed potato lots, both species were frequently detected individually, yet direct co-occurrence of *D. solani* and *P. atrosepticum* was observed in only one sample. Degefu described this near absence of co-occurrence as unlikely to have occurred by chance and proposed that species selection, potentially involving antagonistic interactions, could contribute to this pattern (45). The author highlighted the previously reported ability of *D. solani* to outcompete *Pectobacterium* species, while emphasizing that the mechanisms underlying the apparent exclusion of *D. solani* and *P. atrosepticum* remained unresolved. Our demonstration that *D. solani* efficiently kills *P. atrosepticum* through its T6SS identifies a potential molecular mechanism that could contribute to such competitive interactions. However, whether T6SS-mediated antagonism contributes to the distribution or co-occurrence of these pathogens during potato colonization remains to be established.

Functionally, the ArcZ-SlyA cascade therefore provides a common regulatory route coordinating two mechanistically and spatially distinct forms of microbial antagonism: a contact-dependent antibacterial apparatus and a diffusible antifungal metabolite. Such coordinated regulation could allow *D. solani* to deploy distinct competitive strategies against bacterial and fungal competitors encountered across its ecological niches.

Finally, the phylogenetic distribution of the ArcZ-*slyA* interaction provides a model for regulatory network evolution. While the ArcZ regulatory core is conserved throughout *Enterobacterales* (11), the complementary sequence within the *slyA* mRNA 5′ UTR is restricted to *Pectobacteriaceae* (Fig. 2E; Fig. S3). This distribution is consistent with the emergence of an ArcZ-responsive target site through sequence evolution within the *slyA* mRNA leader, rather than through major changes in the conserved sRNA. The acquisition of a functional sRNA target site within a single 5′ UTR could therefore integrate an existing transcription factor and its downstream regulon into a post-transcriptional program. In this case, a single sRNA-mRNA interaction connects ArcZ to both T6SS-mediated antibacterial competition and solanimycin-dependent antifungal activity. More broadly, our findings illustrate how evolution of a regulatory sequence within a non-coding mRNA region can rewire conserved regulatory circuits and coordinate lineage-specific ecological functions.

## Materials and Methods

A full list of bacterial strains, plasmids, and oligonucleotides used in this study is provided in Tables S4 and S5. Detailed descriptions of all experimental procedures and associated references are provided in SI Appendix, Materials and Methods. These include bacterial growth conditions, strain construction and genetic engineering, Hfq RIL-seq, RNA-seq, quantitative proteomics, purification of His_6_-SlyA, EMSA, RNA isolation, 5’RACE, Northern blot, sequence alignment, prediction of RBS accessibility, interbacterial competition (T6SS killing), yeast growth inhibition assays and statistical analyses. RNA-seq and RIL-seq data have been deposited in the NCBI Gene Expression Omnibus under accession no. GSE348202 and GSE347875, respectively. Mass spectrometry proteomics data have been deposited in the MassIVE repository under accession no. MSV000103229

## Acknowledgments and Funding Sources

We acknowledge the contribution of SFR Biosciences (Université Claude Bernard Lyon 1, CNRS UAR3444, Inserm US8, ENS de Lyon) Protein Science Facility, especially Frédéric Delolme and Adeline Page, for the mass spectrometry analyses. We gratefully acknowledge support from the CNRS/IN2P3 Computing Center (Lyon, France) for providing computing and data-processing resources. This work was supported in part by the Agence Nationale de la Recherche: grant ANR-22-CE35-0017 attributed to E.G., grants ANR-17-CE11-0009 and ANR-25-CE11-6139 attributed to L.A. K.P. acknowledges funding by the DFG (EXC 2051; Project number 390713860 and PA2820/7-1; Project-ID 544846468) and the European Research Council (ArtRNA, CoG-101088027). This work also received support from the INSA Lyon ENJEU program attributed to Q.D.

## Author contributions

Q.D., L.A. and E.G. designed research; Q.D., M.S. and S.K. performed research; Q.D., M.S., S.K., K.P., L.A. and E.G. analyzed data; Q.D., A.R., K.P., L.A. and E.G. provided funding; Q.D., L.A. and E.G. wrote the paper with contribution of all other authors.

