## Supplementary material for "The ArcZ small RNA activates SlyA production to control contact-dependent antibacterial killing and diffusible antifungal activity in *Dickeya solani*": Material and methods appendix

**Materials and Methods**

**Bacterial Strains, Plasmids, and Growth Conditions**

The *Escherichia coli*, *Dickeya solani* and *Pectobacterium atrosepticum* strains and plasmids used in this study are listed in Table S4. Oligonucleotides are listed in Table S5. Kluyveromyces lactis MWL9S1 was used in this study. *E. coli* was routinely grown at 37°C in Luria-Bertani (LB) broth. The yeast *K. lactis* was grown at 30°C in YPG rich medium (1% Bacto yeast extract, 1% Bacto peptone, 2% glucose). *D. solani* and *P. atrosepticum* strains were cultivated in LB unless otherwise specified. For yeast inhibition assays, *D. solani* strains were grown in M63 minimal medium supplemented with 1% sucrose (2 g (NH_4_)_2_SO_4_, 13.6 g KH2PO4, 2.5 mg FeSO_4_ 7H_2_O, 0.2 g MgSO_4_ 7H_2_O, 10 g sucrose, per liter). For growth curve measurements, strains were grown in 200 µL LB in 96-well plates placed inside a temperature-controlled plate reader (Tecan Infinite M200 PRO) with intermittent shaking (100 rpm, 60 s every 15 min) at 30°C, and OD_600_ was monitored every 15 min.

When required, antibiotics were added at the following concentrations: ampicillin (Amp), 100 µg/mL; streptomycin (Sm), 20 µg/mL; gentamicin (Gm), 10 µg/mL and chloramphenicol (Cm), 4 µg/mL, for both *D. solani* and *E. coli*. Diaminopimelic acid (DAP) was supplemented at 57 µg/mL for growth of the *E. coli* MFDpir strain. Solid media contained 12 g/L agar.

**Construction of Gentamicin-marked *P. atrosepticum* Strains**

For type VI secretion system competition experiments, *P. atrosepticum* was marked with a gentamicin resistance cassette by inserting a mini-Tn7-Gm cassette into the attTn7 site using the pTn7-M and pTnS3 plasmids, as previously described for *D. solani* D s0432-1 (1).

**Creation of in-Frame Deletion Mutants**

In-frame deletion mutants were constructed in 2 steps using the *sacB* counter-selection method (2). First, plasmids carrying the upstream and downstream region flanking the region to be deleted and a sacB/cat cassette were constructed. Plasmids pRE112 or pRE112-lacZ𝛼, which are non-replicative in *D. solani*, were used as backbones (3). Once obtained in *E. coli*, the recombinant suicide plasmids were transferred into *D. solani* by conjugation. Recombination allows the plasmid to integrate into the chromosome at either the downstream or upstream target region (selection using chloramphenicol). A second recombination event in the other flanking region generates an in-frame deletion mutant of the target region (selection using sucrose). All oligonucleotides used in this study are listed in Table S5. The *D. solani* Δ*slyA* and Δ*tssM* strains bear a 438-bp and 3498-bp deletion, respectively, centered on the open reading frames of the corresponding genes.

To make the recombinant suicide plasmids, two PCR fragments corresponding to the upstream and downstream approximately 500 bp flanking regions of the target gene or cluster were cloned into pRE112 or pRE112-lacZ𝛼, linearized using the restriction enzymes SacI/KpnI or primers L758/L1227 respectively, using TLTC or Gibson assembly (4, 5). Chemically competent *E. coli* DH5α λpir cells were transformed with the recombinant plasmids using the Mix and Go kit (Zymo Research) and transformants were selected on LB agar supplemented with chloramphenicol. Plasmids were verified by colony PCR with primers L762/L763, restriction digestion, and Sanger sequencing (Eurofins). Chemically competent *E. coli* MFDpir (6) were transformed with the recombinant plasmids and subsequently used as donor strains for conjugation with *D. solani*. For conjugation, equal volumes of *D. solani* and *E. coli* MFDpir cultures were mixed, centrifuged, resuspended in 90 µL LB supplemented with DAP, spotted onto LB agar and incubated at 30°C for 18 h. Bacteria were then resuspended in 1 mL LB, serially diluted, and plated on LB agar supplemented with chloramphenicol to select *D. solani* transconjugants in which the suicide plasmid had integrated into the chromosome by a first homologous recombination event. Transconjugants were subsequently plated on LB agar without NaCl supplemented with 5% sucrose and incubated at 19°C for 2 to 3 days to select for a second homologous recombination event resulting in plasmid excision. Sucrose-resistant colonies were patched on LB agar supplemented with chloramphenicol to verify plasmid loss. All deletions were confirmed by colony PCR using flanking primers listed in Table S5.

**Construction of Dual Reporter Fusions *slyA*-*sfGFP*_*mCherry* and *slyA**-*sfGFP*_*mCherry***

Translational and transcriptional reporter fusions were integrated at the native *slyA* locus using the same allelic-exchange procedure described above. To build the reporter cassette, *slyA* was PCR-amplified together with its promoter and 5′ UTR but without its stop codon, and fused in frame to *sfGFP*; a *mCherry* gene preceded by a strong synthetic RBS was placed immediately downstream on the same transcript. In this dual reporter, the SlyA‑sfGFP fusion protein is translated from the native *slyA* start and RBS and therefore reports SlyA translation, whereas mCherry is translated from an independent RBS on the same mRNA and serves as a proxy for transcript level. For this study, the mCherry readout was not used. The upstream and downstream ~500-bp homology regions flanking the insertion site were amplified and assembled with the *slyA*-*sfGFP* and RBS_*mCherry* fragments into pRE112-lacZα. Chemically competent *E. coli* DH5α λpir cells were transformed with the construct. Transformants were selected on LB agar supplemented with chloramphenicol, verified by colony PCR with primers L762/L763, restriction digestion, and Sanger sequencing (Eurofins). Chemically competent *E. coli* MFDpir cells were transformed with the recombinant plasmid and subsequently used as donor cells for conjugation with *D. solani*. Allelic exchange was performed as described above for the construction of in-frame deletion mutants. For translational fusion assay, bacterial cultures were inoculated at OD_600_ = 0.006 in 96-well plates in LB. The cultures were incubated at 30°C and GFP was measured and OD_600_ were measured every 10 minutes using a Tecan Infinite M200 plate reader. GFP values were normalized to OD_600_. Four biological replicates were performed.

**Construction of Promoter Fusions and Luminescence Assays**

To monitor promoter activity of *hcpA*, *hcpB*, *tssB*, *vgrGB*, the promoter regions of these genes from the *D. solani* strain D s0432-1 were amplified by PCR using primer pairs listed in Table S5. These 300‑bp promoter fragments (P_xxx_) were cloned into pSEVA421-P*_solA_*_*luxCDABE-gfp* (1) linearized with primers L2590/L2591 by HiFi assembly. Chemically competent *E. coli* DH5α λpir cells were transformed with the resulting pSEVA421-Pxxx-luxCDABE-gfp plasmids. Transformants were selected on LB agar supplemented with streptomycin (Sm), and plasmids were verified by colony PCR with primers L707/L33, restriction digestion, and Sanger sequencing (Eurofins). Chemically competent *E. coli* MFDpir cells were then transformed with the verified plasmids and used as donor strains for conjugation with *D. solani*. For luminescence assays, bacterial cultures were inoculated at OD_600_ = 0.006 in 96-well plates in LB for P*hcpA*, P*hcpB*, P*tssB* and P*vgrGB*, and in M63 medium supplemented with 1% sucrose for P*solA*. The cultures were incubated at 30°C and luminescence was measured at OD_600_ = 0.3 using a Tecan Infinite M200 plate reader with an integration time of 1 s per well. Luminescence values were normalized to OD_600_. Four biological replicates were performed.

**Construction of the *slyA* and *tssM* Complementation Plasmids.**

The *slyA*, and *tssM* genes under the control of their own promoter were amplified by PCR from the *D. solani* strain D s0432‑1 with oligonucleotide pairs L1997/ L1998, and L2693/ L2694, respectively. These fragments were cloned into pWSK29-oriT (1) linearized with primers L2634/L2635 or using the restriction enzyme EcorV. Chemically competent *E. coli* DH5α λpir cells were transformed with the resulting constructs. Transformants were selected on LB agar supplemented with ampicillin, verified by colony PCR, restriction digestion, and Sanger sequencing (Eurofins). Chemically competent *E. coli* MFDpir cells were transformed with the verified plasmids and subsequently used as donor strains for conjugation with *D. solani*. Transconjugants were selected on LB agar supplemented with ampicillin.

**Hfq RIL-seq and Computational Analysis**

To construct the *D. solani* D s0432-1 strain expressing C-terminally 3×FLAG-tagged Hfq, the ~500-bp upstream and downstream homology regions flanking *hfq* were amplified using primers designed to introduce the 3×FLAG sequence at the C terminus of Hfq and assembled into pRE112-*lacZα* (Table S5). Chemically competent *E. coli* DH5α λpir cells were transformed with the resulting recombinant plasmid. Transformants were selected on LB agar supplemented with chloramphenicol, verified by colony PCR with primers L762/L763, restriction digestion, and Sanger sequencing (Eurofins). Chemically competent *E. coli* MFDpir cells were transformed with the recombinant plasmid and subsequently used as donor cells for conjugation with *D. solani*. Allelic exchange was performed as described above for the construction of in-frame deletion mutants. Allele replacements were confirmed by colony PCR using flanking primers listed in Table S5.

RIL-seq was performed following the original protocol (7). Briefly, a *D. solani* D s0432-1 strain expressing a chromosomal C-terminal 3×FLAG-tagged Hfq was grown in LB at 30°C to low cell density (LCD, OD_600_ = 0.5) and high cell density (HCD, OD_600_ = 1.0), with three biological replicates per condition. An isogenic strain expressing untagged Hfq was processed in parallel at both cell densities as a negative control for non-specific RNA recovery. At OD_600_ = 0.5 and OD_600_ = 1, transcription was stopped by addition of 0.5 volumes of cold 100% methanol. Cells corresponding to 50 OD_600_ units were subjected to UV crosslinking of protein-RNA complexes, followed by lysis and co-immunoprecipitation with a monoclonal anti-FLAG antibody (Sigma; F1804). Co-immunoprecipitated RNA was trimmed with RNase A/T1, and proximal RNA ends were ligated with T4 RNA ligase. After proteinase K digestion, RNA was extracted, fragmented, and treated with TurboDNase. Ribosomal RNA was depleted using biotinylated rRNA-specific probes: co-immunoprecipitated RNA was combined with 1× SSC, 1 mM EDTA and a biotinylated oligonucleotide mix (5.8 nM for each probe: 16S, 23S, 5S oligos), denatured, cooled, and incubated with streptavidin beads (ThermoFisher; 65001) for 5 min at room temperature followed by 5 min at 50 °C (8). rRNA-depleted RNA was purified with the Agencourt AMPure XP kit (Beckman Coulter) and used for cDNA library preparation; libraries were amplified with NEBNext Ultra II Q5 polymerase and sequenced on a NextSeq 1000 system in paired-end mode (200-nt read length). RNA recovered from *D. solani* consistently displayed relatively low integrity, a recurrent feature in our preparations. Similar difficulties in recovering high-integrity RNA have previously been reported for the closely related species *D. dadantii* (9), suggesting that this may represent a technical limitation associated with RNA isolation from *Dickeya spp*. Nevertheless, all biological replicates yielded sufficient material for cDNA library preparation and sequencing.

Computational analysis was performed with ChimericFragments (10). Demultiplexed reads were quality- and complexity-filtered and mapped to a *D. solani* reference genome derived from the IPO 2222 assembly (NCBI accession CP015137.1). Whole-genome comparative analysis confirmed that the D. solani D s0432-1 and IPO 2222 genomes share >99.99% Average Nucleotide Identity (ANI) with complete collinear synteny and 100% orthologous gene conservation, ensuring comprehensive and unbiased read mapping. Chimeric reads representing RNA-RNA interactions were identified and annotated (ncRNA and UTR_CDS), and interactions were filtered using a cut-off of 5 reads per interaction and an base paired-FDR-adjusted p‑value ≤ 0.05. Data analysis and visualization were performed according to the previously published computational pipeline from Papenfort Lab (10). Raw reads and RNA-RNA interaction data of the RIL-Seq experiments were deposited into the Gene Expression Omnibus (GEO) repository under accession number GSE347875.

**Global Gene Expression Profiling by RNA-Seq**

*D. solani* strains D s0432-1 WT (DS49) and Δ*slyA* (DS753) were cultivated in triplicate in LB medium at 30°C with 120 rpm shaking. LB medium was selected because sufficient RNA integrity for library preparation and sequencing could not be achieved when strains were cultivated in M63 minimal medium supplemented with 1% sucrose. At OD_600_ = 0.5, transcription was stopped by addition of 0.5 volumes of cold 100% methanol. Total RNA was isolated by hot phenol extraction, treated with Turbo DNase (Thermo Fisher Scientific), and RNA integrity was confirmed using a Bioanalyzer (Agilent). Ribosomal RNA was depleted using rRNA-specific biotinylated probes as previously described (8). rRNA-depleted RNA was purified using Agencourt AMPure XP beads (Beckman Coulter) and fragmented for 5 min at 75°C using the NEBNext Magnesium RNA Fragmentation Module (NEB). cDNA libraries were prepared using the NEBNext UltraExpress RNA Library Prep Kit for Illumina (NEB, E3330L) according to the manufacturer's instructions. Library quality was assessed on an Agilent 2100 Bioanalyzer and pooled libraries were sequenced on a NextSeq 1000 system with 50 nt paired‑end sequencing mode. Demultiplexed reads were trimmed and mapped to the *D. solani* type strain IPO 2222 reference genome (NCBI accession CP015137.1) using the RNA-Seq Analysis tool of CLC Genomics Workbench (Qiagen) with standard parameters. Differential expression analysis was performed using the DESeq2 R package. Differentially expressed genes were defined as those with a log_2_ fold change ≤‑1 or ≥ 1 and an FDR‑adjusted p‑value ≤ 0.05. Three biological replicates were performed per strain. The RNA-seq data have been deposited in the NCBI Gene Expression Omnibus (GEO) repository under accession number GSE348202.

**Quantitative Proteomics**

Quantitative proteomics was performed on the *D. solani* D s0432-1 Δ*slyA* deletion mutant relative to the WT strain to provide an independent line of evidence for the ArcZ‑dependent expression program at the protein level. M63 minimal medium supplemented with 1% sucrose was used because it supports robust protein extraction and reproducible quantification under defined nutritional conditions.

*D. solani* strains DS49 (WT) and DS753 (Δ*slyA*) were cultivated in triplicate in M63 minimal medium supplemented with 1% sucrose at 30°C with 120 rpm shaking to OD_600_ = 0.3. Cells were harvested by centrifugation and protein extracts were prepared with the EasyPepTM mini kit (ref. A40006, ThermoFisher Scientific) according to the manufacturer protocol. Briefly, proteins were denaturated, reduced and alkylated (10 min, 95°C, 1000 rpm), then digested with a mixture of endoproteinase Lys-C/trypsin (3h, 37°C, 500 rpm). After a cleaning step, peptides were dried, suspended in 0.1% formic acid and dosed with the quantitative fluorometric peptide assay (ref. 23290, Thermo Scientific).

200 ng of each sample were analyzed on the Exploris 480 mass spectrometer coupled with a Vanquish NEO nanoLC system (Thermo Scientific). Peptides samples were loaded on a C18 Acclaim PepMap100 trap-column 300 µm ID x 5 mm, 5 µm, 100Å (ThermoFisher Scientific) and separated on a C18 Acclaim Pepmap100 nano-column, 50 cm x 75 µm i.d, 2 µm, 100 Å (Thermo Scientific) with a 45 minutes linear gradient from 3% to 25% buffer B (A: 0.1% FA in H2O, B: 0.1% FA in ACN/H2O (80/20)) from 25% to 35% of B in 15 min and then from 35% to 100% of B in 0.1 min, hold for 12 min and followed by a washing and equilibration steps for a total duration of 72 minutes. The flow rate was 300 nL/min and the column temperature was kept constant at 45°C. Peptides were analysed with a DDA 1s HCD method: MS data were acquired in a data dependent strategy (DDA) selecting the fragmentation events based on the most abundant precursor ions in a 1s survey scan (350-1400 Th). Resolutions of the survey and MS/MS scans were respectively set at 120,000 and 15,000 at m/z 200 Th. The Ion Target Values for the survey and the MS/MS scans in the Orbitrap were set to 3E6 (300%) and 1E5 (100%) respectively and the maximum injection time was set to 50 ms for MS scan and 22 ms for MS/MS scan. Parameters for acquiring HCD MS/MS spectra were as follows: collision energy = 30 and isolation window = 4 m/z. The precursors with unknown charge state, charge state of 1 and 6 or greater than 6 were excluded. Peptides selected for MS/MS acquisition were then placed on an exclusion list for 40s using the dynamic exclusion mode to limit duplicate spectra.

Raw data were processed with Proteome Discoverer 3.1 (Thermo Scientific) through the Chimerys and Sequest HT search engines against uniparc *D. solani* D s0432-1 (4951 entries, version june 2024) and a contaminant database. Mass tolerance was set at 20 ppm, and 2 missed cleavage were allowed. Oxidation (M), acetylation (N-ter protein) were set as variable modifications and Carbamidomethylation (C) as fixed modification. Validation of identified peptides and proteins was done using a target decoy approach with a false discovery rate positive (FDR < 1%). Protein quantitation was performed with precursor ions quantifier node in Proteome Discoverer 3.1 software. Proteins were normalized to the total peptide amount and ratios of quantitation were calculated in a pairwise way. Statistical validation was based on t-test (a protein is considered differentially expressed between two conditions if the log_2_ fold change is ≤-1 or ≥ 1 and have a p-value < 0.05). Proteomics analyses were performed at the Protein Science Facility of SFR Biosciences (UAR3444/CNRS, US8/Inserm, ENS de Lyon, UCBL), Lyon, France. The mass spectrometry proteomics data have been deposited to the Center for Computational Mass Spectrometry repository (University of California, San Diego) via the MassIVE tool with the dataset identifier MassIVE MSV000103229.

**Purification of His_6_-SlyA**

Construction of the pIBA37-6×His-*slyA* expression vector. A codon-optimized *slyA* fragment was synthesized by Twist Bioscience. The synthetic fragment was first cloned into pGEX-6P-3 digested with BamHI and XhoI. The resulting pGEX-6P-3-*slyA* construct was verified by restriction analysis and Sanger sequencing. *slyA* was then amplified from pGEX-6P-3-*slyA* using primers L2177 and L2178 and cloned into BsaI-linearized pIBA37, generating the pIBA37-6×His-*slyA* expression plasmid.

A N-terminal His_6_-tagged SlyA (His_6_-SlyA) was expressed from pIBA37-Nter-His_6_-*slyA* in *E. coli* BL21. A fresh transformant was used to inoculate an overnight LB-ampicillin preculture; 5 mL were then used to inoculate 100 mL LB‑ampicillin. Expression was induced at OD_600_ = 0.4 with 200 ng/mL anhydrotetracycline and continued for 3 to 4 h at 37°C. Cells were harvested by centrifugation (10,000 × g, 10 min) and stored at ‑80°C.

Purification was performed under native conditions by Ni‑NTA batch chromatography, adapted from a standard protocol (QIAGEN). The pellet was thawed on ice, resuspended in 10 mL lysis buffer NPI‑10 (50 mM NaH_2_PO_4_, 300 mM NaCl, 10 mM imidazole, pH 8.0) supplemented with 1 mg/mL lysozyme, and incubated on ice for 30 min. The lysate was sonicated on ice until clarified and cleared by centrifugation (10,000 × g, 30 min, 4°C). The soluble fraction was incubated with 100 µL Ni‑NTA resin; the resin was washed with wash buffer NPI‑20 (50 mM NaH_2_PO_4_, 300 mM NaCl, 20 mM imidazole, pH 8.0) and the protein eluted with elution buffer NPI‑500 (50 mM NaH_2_PO_4_, 300 mM NaCl, 500 mM imidazole, pH 8.0) in a final volume of 500 µL. The eluate was buffer-exchanged and concentrated on a Vivaspin 500 device (10 kDa MWCO, 12,000 × g) into storage buffer (20 mM Tris-HCl pH 8.0, 100 mM NaCl, 20% glycerol). Protein purity was verified by SDS-PAGE (Fig. S5B) and concentration determined by Bradford and BCA assays. Throughout, SlyA concentrations are expressed as monomer.

**Electrophoretic Mobility Shift Assay (EMSA)**

Promoter DNA probes (P*hcpA: 158 bp,* P*hcpB: 171 bp,* P*solA*: 150 bp and negative fragment (*slyA* CDS): 162 bp) were generated by PCR using a 5′‑6‑FAM‑labelled forward primer (Table S5) and purified with Mag‑Bind TotalPure NGS magnetic beads (Omega Bio‑tek). Binding reactions (10 µL final volume) contained 10 nM labelled probe and purified His_6_-SlyA monomer at 0, 100, 200, 300, 400, 500, 600 and 700 nM in binding buffer (10 mM Tris‑HCl pH 8.0, 50 mM KCl, 1 mM EDTA, 0.1 mM DTT, 10 µg/mL BSA, 10% glycerol). Reactions were incubated at 30°C for 30 min. A FAM‑labelled fragment internal to the *slyA* coding sequence was used as a non‑specific negative‑control probe. Polyacrylamide 8% gels (37.5:1) were pre‑run in 0.5x TBE at 50 V for 20 min. Then samples were loaded and migrated at 90 V for 90 min at 4°C in 0.5x TBE. Labelled DNA was visualized on a ChemiDoc imaging system (Bio-Rad) using excitation at 480 nm and emission at 520 nm. Band intensities were quantified using Vilber software. The bound fraction (shifted signal divided by total signal) was plotted against SlyA concentration to estimate apparent half-maximal binding concentrations (Fig. 4C). Two independent experiments were performed.

**5′ RACE mapping of the *slyA* transcription start site**

The transcription start site of *slyA* was determined by template-switching 5′ rapid amplification of cDNA ends (5′ RACE) using the Template Switching RT Enzyme Mix (New England Biolabs, NEB #M0466), following the manufacturer’s protocol. Total RNA was annealed to L2109 a *slyA*-specific reverse-transcription primer together with dNTPs for 5 min at 70°C. Reverse transcription and template switching were then performed in a 10-µl reaction containing 1× Template Switching RT Buffer, 3.75 µM template-switching oligonucleotide (TSO), and 1× Template Switching RT Enzyme Mix. Reactions were incubated for 90 min at 42°C, followed by enzyme inactivation for 5 min at 85°C.

The resulting cDNA was diluted twofold and the 5′ region of *slyA* was amplified using Q5 Hot Start High-Fidelity 2× Master Mix (NEB #M0494), a TSO-specific forward primer (L2152), and a *slyA*-specific reverse primer (L2110). PCR products were directly subjected to Sanger sequencing. The *slyA* transcription start site was assigned to the first genomic nucleotide immediately adjacent to the TSO-derived sequence after alignment of the obtained sequence to the *D. solani* genome.

**RNA Isolation and Northern Blot Analysis**

For ArcZ quantification, *D. solani* D s0432-1 strain was grown at 30°C with  120 rpm shaking in LB medium to OD_600_ = 0.5; 0.8; and 1. One‑milliliter aliquots were fixed by mixing with 1 mL cold methanol (4°C) and harvested by centrifugation. Cell pellets were stored at ‑80°C prior to extraction. RNA was extracted using the RNAsnap method(11). Pellets were resuspended in 100 µL RNAsnap buffer (18 mM EDTA pH 8.0, 0.025% SDS, 95% formamide) and heated twice at 95°C for 7 min. The lysate was mixed with  650 µL Tri-Reagent (Zymo Research), and RNA was purified using the Direct‑zol RNA MiniPrep kit (Zymo Research) according to the manufacturer's instructions. RNA was eluted in 30 µL DNase/RNase-free water and quantified by NanoDrop spectrophotometry.

For Northern blot analysis, 1.2 µg of total RNA was resolved per lane on an 8% acrylamide‑urea gel (Invitrogen) with 1 µL Low Range ssRNA Ladder (New England Biolabs) as a size standard. Electrophoresis was performed at 200 V in TBE buffer. RNA was transferred to a nylon membrane (Hybond‑N+, GE Healthcare) by electrophoretic transfer in 0.5x TBE (30 min at 300 mA, Transblot, Bio-Rad) and crosslinked by UV‑irradiation ( 254‑nm, Stratalinker, Stratagene). Membranes were pre‑hybridized for 4 h at 42°C in ULTRAhyb buffer (Ambion) and hybridized overnight at 42°C with the biotinylated probe LA196 (5'-Biotin-

AAAAAAAATGACCCCGACCGAGGTCGGGGTGCGCGAATTATGCGCCAACACCAGGGAAAGCG-

TgBiotin-3', purchased from Sigma-Aldrich). Membranes were washed at 65°C with 2x SSC containing 0.1% SDS. Hybridized probes were detected using the Chemiluminescent Nucleic Acid Detection Module (Pierce) with streptavidin-HRP and luminol. Chemiluminescence signals were acquired using a CCD camera imaging workstation (Thermo Scientific).

**Nucleotide Sequence Alignment, Prediction of RBS Accessibility and IntaRNA Interaction Prediction**

To assess the conservation of the ArcZ-*slyA* interaction across *Enterobacterales*, comparative sequence analysis was performed across the *Pectobacteriaceae* and against representative non‑*Pectobacteriaceae* taxa. The *slyA* 5′ UTR sequences, defined as the 100 nt region upstream of the *slyA* start codon, were retrieved from the NCBI nucleotide database (https://blast.ncbi.nlm.nih.gov) for the 18 species listed in Fig. 2E, Fig. S3A and Fig. S3B. Multiple sequence alignments were performed using ClustalW (12) via the DDBJ ClustalW web server (https://www.genome.jp/tools-bin/clustalw).

Alignment outputs were visualized and annotated using Jalview version 2.11.5.1 (13).

The accessibility of the *slyA* ribosome-binding site (RBS) was estimated with the ViennaRNA package (version 2.7.2, Python interface) (14). The *slyA* mRNA fragment used for the analysis extended from 94 nt upstream to 15 nt downstream of the translation start codon. Base-pairing probabilities were calculated from the partition function at 37°C, and the probability of each nucleotide being single-stranded was defined as one minus the summed probability of its pairing with any other nucleotide. RBS accessibility was calculated as the mean single-stranded probability across the region spanning the Shine-Dalgarno sequence and the start codon (positions −10 to +3 relative to the A of the AUG start codon). To assess the effect of ArcZ binding, ArcZ and the *slyA* mRNA fragment were co-folded using the ViennaRNA dimer partition function, and RBS accessibility was recalculated within the resulting ArcZ-*slyA* complex. These values represent thermodynamic predictions and do not account for Hfq-dependent effects.

Interactions between ArcZ and *slyA* mRNA were predicted with IntaRNA (version 3.4.1; Freiburg RNA Tools web server) (15), using 3′‑processed ArcZ as the query and the *slyA* fragment 145 nt before the start codon to 15 nt after the start codon (which contains the 5′ UTR ) as the target, with default parameters. The predicted hybridization free energy (ΔG) and interaction site were recorded for each species (Fig. S3A).

**RNA fold and AlphaFold-based prediction of the ArcZ-*slyA* mRNA interaction**

Secondary-structure predictions were performed using the ViennaRNA package 2.0. A part of *slyA* 5′ UTR fragment adjacent to the RBS (5′-AUUGCAAUCCAGGGAAGUUGCUUAGUAUGCUAACUAUAAGGAGAAGUGAUG-3′) was folded alone using RNAfold. To model the ArcZ‑*slyA* complex, the same *slyA* fragment was co-folded with processed ArcZ sequence (5′-UUUCCCUGGUGUUGGCGCAUAAUUCGCGCACCCCGACCUCGGUCGGGGUCAUUUUUUUU-3′) using RNAcofold. Secondary structures were visualized using VARNA (16).

Three-dimensional RNA structures were predicted using the AlphaFold Server, powered by AlphaFold 3 (17) . Predictions were performed for the *slyA* 5′ untranslated region (5′ UTR ) alone and for the ArcZ-*slyA* mRNA complex with the same RNA input used for secondary structure prediction. For prediction of the *slyA* 5′ UTR alone, the same *slyA* RNA sequence was submitted in the absence of ArcZ.

Predictions were generated using the default AlphaFold Server parameters. For each prediction, the highest-ranked model according to the AlphaFold confidence/ranking score was selected for subsequent structural inspection and comparison. Predicted structures were visualized using Mol* Viewer (18).

**Interbacterial Competition (T6SS killing) Assays**

Interbacterial competition assays were performed using *D. solani* as the attacker and a Gentamicin‑resistant (Gm^R^) *P. atrosepticum* SCRI1043 strain as the prey. Attacker and prey strains were grown overnight in LB, then diluted to OD_600_ = 0.05 and grown to OD_600_ = 0.5 at 30°C. Attacker and prey were mixed at a 10:1 (attacker: prey) ratio and harvested by centrifugation (4,000 × g, 5 min). The supernatant was removed and cells were re-pelleted resuspended in 15 µL LB. A 5 µL drop of the mixture was spotted onto a 0.45 µm membrane filter placed on an LB agar plate, and a parallel 5 µL aliquot was serially diluted and plated (on LB agar plates without antibiotics, and with gentamycin) to determine the initial attacker‑to‑prey ratio (Fig. S6A). After 16 h of incubation at 30°C, the filter was recovered from the plate, resuspended in 1 mL LB, serially diluted, and plated on LB and on LB supplemented with gentamicin (10 µg/mL) to determine the attacker‑to‑prey ratio post competition. As *D. solani* is gentamicin‑sensitive, prey survival was quantified as the log_10_ CFU of *P. atrosepticum* recovered on gentamicin-containing plates. Four biological replicates were performed.

**Yeast Growth Inhibition Assay**

YPG agar medium was melted, cooled to approximately 45°C, and mixed with a *K. lactis* overnight culture at 2 OD_600_ units per 30 mL agar to prepare seeded plates. Strains *D. solani* D s0432-1 (WT) and mutants were grown at 30°C with  120 rpm shaking in M63 minimal medium supplemented with 1% sucrose until they reached OD_600_ = 2. Five microliters of each *D. solani* culture were spotted onto *K. lactis* seeded plates and incubated at 30°C for 24 to 48 h prior to visualization of inhibition zones. Four biological replicates were performed for each tested strain.

**Statistical Analysis**

All statistical analyses were performed using GraphPad Prism version 10. Comparisons between two groups were performed using the Mann-Whitney test. For growth curves, doubling times were calculated during the exponential growth phase and compared between strains using pairwise Mann‑Whitney tests. For proteomic data, differential protein abundance was assessed by t-test with a p‑value threshold of 0.05, as described above. For transcriptomic data, differential gene expression was assessed using DESeq2 with median-of-ratios normalization and an FDR-adjusted p‑value threshold of 0.05, as described above.

Results are expressed as mean ± standard deviation (SD) unless otherwise stated. Significance thresholds were set at p < 0.05 (*), p < 0.01 (**), and p < 0.001 (***). Non‑significant differences are indicated as ns.

10.	Siemers M, Lippegaus A, Papenfort K. 2024. ChimericFragments: computation, analysis and visualization of global RNA networks. NAR Genom Bioinform 6:lqae035.

17.	Abramson J, Adler J, Dunger J, Evans R, Green T, Pritzel A, Ronneberger O, Willmore L, Ballard AJ, Bambrick J, Bodenstein SW, Evans DA, Hung C-C, O’Neill M, Reiman D, Tunyasuvunakool K, Wu Z, Žemgulytė A, Arvaniti E, Beattie C, Bertolli O, Bridgland A, Cherepanov A, Congreve M, Cowen-Rivers AI, Cowie A, Figurnov M, Fuchs FB, Gladman H, Jain R, Khan YA, Low CMR, Perlin K, Potapenko A, Savy P, Singh S, Stecula A, Thillaisundaram A, Tong C, Yakneen S, Zhong ED, Zielinski M, Žídek A, Bapst V, Kohli P, Jaderberg M, Hassabis D, Jumper JM. 2024. Accurate structure prediction of biomolecular interactions with AlphaFold 3. Nature 630:493–500.

18.	Sehnal D, Bittrich S, Deshpande M, Svobodová R, Berka K, Bazgier V, Velankar S, Burley SK, Koča J, Rose AS. 2021. Mol* Viewer: modern web app for 3D visualization and analysis of large biomolecular structures. Nucleic Acids Res 49:W431–W437.
