## Supplementary material for "The ArcZ small RNA activates SlyA production to control contact-dependent antibacterial killing and diffusible antifungal activity in *Dickeya solani*": S4 Table

**Table S4. Bacterial strains and plasmids used in this study.**

All *Dickeya solani,* *Escherichia coli* and *Pectobacterium atrosepticum* strains and plasmids used in this study are listed with their genotype, relevant genetic modifications, antibiotic-resistance markers and source or reference.

**Bacterial strain and plasmid**

| **Strains** | Description | Source |
| --- | --- | --- |
| ***Escherichia coli* K12** |  |  |
| DH5α | supE44 lacU169 (Φ80lacZ∆ M15) hsdR17 (rK mK ) recA1 endA1 gyrA96 thi-1 relA1 | Laboratory collection |
| DH5α λpir | λpir phage lysogen of DH5α | Laboratory collection |
| MFDpir | RP4-2-Tc::(∆Mu1::aac(3)IV-∆aphA-·∆nic35-∆Mu2::zeo) ∆dapA::erm-pir) ∆recA | (1) |
| BL21 | F⁻, ompT hsdSB(rB⁻ mB⁻) gal dcm | Laboratory collection |

***Dickeya solani* D s0432-1**

| DS49 | *D. solani* D s0432-1 WT *arcZ_1_* | Laboratory collection |
| --- | --- | --- |
| DS354 | DS49 *D. solani* D s0432-1 ∆*arcZ_1_* | (2) |
| DS433 | DS49 *D. solani* D s0432-1 pEGL332 | (2) |
| DS439 | DS354 *D. solani* D s0432-1 ∆arcZ*_1_* /pEGL332 | (2) |
| DS441 | DS354 *D. solani* D s0432-1 ∆arcZ*_1_* /pEGL334 | (2) |
| DS1058 | DS49 *D. solani* ∆*solABCDEFGHIJKL*(named ∆*sol*) | (3) |
| DS753 | DS49 *D. solani* D s0432-1 ∆*slyA* | This study |
| DS789 | DS753 *D. solani* D s0432-1 ∆slyA /pEGL332 | This study |
| DS785 | DS753 *D. solani* D s0432-1 ∆*slyA*/pEGL479 | This study |
| DS1056 | DS49 *D. solani* D s0432-1 ∆*tssM* | This study |
| DS1074 | DS1056 /pEGL332 | This study |
| DS1075 | DS1056 /pEGL601 | This study |
| DS909 | DS354 *slyA-*sfGFP_mcherry | This study |
| DS910 | DS49 *slyA-*sfGFP_mcherry | This study |
| DS938 | DS909 /pEGL332 | This study |
| DS939 | DS909 /pEGL334 | This study |
| DS940 | DS910 /pEGL332 | This study |
| DS974 | DS910 *D. solani* D s0432-1 *slyA**(CCCA)-sfGFP_mcherry | This study |
| DS1080 | DS974 /pEGL332 | This study |
| DS1081 | DS974 /pEGL334 | This study |
| DS1015 | DS974 /pEGL584 | This study |
| DS956 | DS49 D. solani D s0432-1 *hfq*::3xFlag | This study |
| DS820 | DS433 /pEGL385/pEGL332 | This study |
| DS821 | DS439 /pEGL385/pEGL332 | This study |
| DS822 | DS789 /pEGL385/pEGL332 | This study |
| DS823 | DS785 /pEGL385/pEGL479 | This study |
| DS1084 | DS433 /pEGL602/pEGL332 | This study |
| DS1085 | DS433 /pEGL605/pEGL332 | This study |
| DS1086 | DS433 /pEGL603/pEGL332 | This study |
| DS1087 | DS433 /pEGL604/pEGL332 | This study |
| DS1088 | DS439 /pEGL602/pEGL332 | This study |
| DS1089 | DS439 /pEGL605/pEGL332 | This study |
| DS1090 | DS439 /pEGL603/pEGL332 | This study |
| DS1091 | DS439 /pEGL604/pEGL332 | This study |
| DS1096 | DS789 /pEGL602/pEGL332 | This study |
| DS1097 | DS789 /pEGL605/pEGL332 | This study |
| DS1098 | DS789 /pEGL603/pEGL332 | This study |
| DS1099 | DS789 /pEGL604/pEGL332 | This study |
| DS1100 | DS785 /pEGL602/pEGL479 | This study |
| DS1101 | DS785 /pEGL605/pEGL479 | This study |
| DS1102 | DS785 /pEGL603/pEGL479 | This study |
| DS1103 | DS785 /pEGL604/pEGL479 | This study |

| ***Pectobacterium atrosepticum*** |  |  |
| --- | --- | --- |
| P859 | *P. atrosepticum* SCRI1043 WT | Laboratory collection |
| P1023 | *P. atrosepticum* SCRI1043 WT *glmS::Tn7-gent*, Gm^R^ | This study |

**Plasmid**

| pRE112 | Suicide vector for allelic exchange, Cm^R^, *sacB*, *oriT* RP4, *ori*R6K | (4) |
| --- | --- | --- |
| pTn7-M | Km^R^ Gm^R^, ori R6K, Tn7L and Tn7R extremities, standard multiple cloning site, oriT RP4 | (5) |
| pTNS3 | Ap^R^, ori R6K, TnsABCD operon, oriT RP4 | (6) |
| pEGL473 | pRE112-*lacZ*𝛼 | (3) |
| pEGL442 | pRE112-∆*slyA* | This study |
| pEGL513 | pRE112-lacZ𝛼-*slyA*-sfGFP_mcherry | This study |
| pEGL555 | pRE112-lacZ𝛼-*hfq*::C-terminal-3xFlag | This study |
| pEGL565 | pRE112-lacZ𝛼-*slyA**(CCCA)-sfGFP_mcherry | This study |
| pEGL596 | pRE112-∆*tssM* | This study |
| pGEX-6P-3 | Expression vector, Amp^R^, IPTG-inductible, N-terminal-GST, ColE1 | (7) |
| pEGL491 | pGEX-6P-3-N-terminal GST-*slyA* optimized | This study |
| pASK-IBA37(+) (pIBA37) | Expression vector, Amp^R^, AHT-inducible, N-terminal 6xHis tag ColE1 | IBA technology |
| pEGL507 | pIBA37-N-terminal-6xHis-*slyA* | This study |
| pWSK29 | Amp^R^, pSC101 ori, lacZp expression vector | (8) |
| pEGL332 | pWSK29-oriT | (2) |
| pEGL334 | pWSK29-oriT-*arcZ_1_* | (2) |
| pEGL479 | pWSK29-oriT-*slyA* | This study |
| pEGL584 | pWSK29-oriT-*arcZ**(TGGG) | This study |
| pEGL601 | pWSK29-oriT-*tssM* | This study |
| pSEVA421 | Sm^R^, *ori RK2, oriT* | (9) |
| pEGL385 | pSEVA421-P*_solA__gfp_luxCDABE* | (2) |
| pEGL602 | pSEVA421-P*_hcpA__gfp_luxCDABE* | This study |
| pEGL603 | pSEVA421-P*_tssB__gfp_luxCDABE* | This study |
| pEGL604 | pSEVA421-P*_vgrGB__gfp_luxCDABE* | This study |
| pEGL605 | pSEVA421-P*_hcpB__gfp_luxCDABE* | This study |
