## Supplementary material for "The ArcZ small RNA activates SlyA production to control contact-dependent antibacterial killing and diffusible antifungal activity in *Dickeya solani*": S5 Table

**Table S5: Oligonucleotides used in this study**

| Oligonucleotide | | Sequence (5’-3’) | | Use |
| --- | --- | --- | --- | --- |
| **L1896** | ACTGCATGaattcccgggagagctcattgcgcgctaacgttacgg | | PCR to amplify upstream fragment of slyA to clone in pRE112 | |
| **L1897** | taattaagactgcaattccatcacttctccttatag | |  |  |
| **L1898** | gtgatggaattgcagtcttaattaatagaaaagtaaatgtcttg | | PCR to amplify downstream fragment of slyA to clone in pRE112 | |
| **L1899** | atcccaagcttcttctagaggtaccgcgttgccggtaacagcattg | |  |  |
| **L2203** | cgccagggttttcccagtcacgtccaggtacgtttgctgaaaaaaag | | PCR to amplify upstream fragment of slyA + CDS slyA without stop codon to clone in pRE112-lacZα | |
| **L2204** | tgaacagttcttcgcctttacgagactggctctcatgtaatgc | |  |  |
| **L1486** | cgtaaaggcgaagaactgttc | | PCR to amplify sfGFP fragment without ATG to clone in pRE112-lacZα to make pRE112-slyA-sfGFP_RBS-mcherry | |
| **L2202** | ttacttatacagctcgtccataccg | |  |  |
| **L2205** | tggacgagctgtataagtaaGAATTCaaagaggagaaatactagatggt | | PCR to amplify mcherry fragment with RBS to clone in pRE112-lacZα to make pRE112-slyA-sfGFP_RBS-mcherry | |
| **L2206** | ttttctattaaattacttgtacagctcgtccat | |  |  |
| **L2207** | gtacaagtaatttaatagaaaagtaaatgtcttggcgg | | PCR to amplify downstream fragment of slyA to clone in pRE112-lacZα to make pRE112-slyA-sfGFP_RBS-mcherry | |
| **L2208** | ATCCATGGTCGACAAGCTTCTAggtcggtaataccgtcgg | |  |  |
| **L2279** | acgccagggttttcccagtcacgtcagtttgcgattgctccgg | | PCR to amplify upstream fragment of hfq with C-terminal 3xFlag to clone in pRE112-lacZα | |
| **L2566** | ttgtagtcgatatcatgatctttataatcaccgtcatggtctttgtagtcttcagcgtcatcactttcctg | |  |  |
| **L2282** | cgaCTCTAGAGGATCCCCGGGTActgagcgaagatatccagaatcaacc | | PCR to amplify downstream fragment of hfq with C-terminal 3xFlag to clone in pRE112-lacZα | |
| **L2567** | TGATTATAAAGATCATGATATCGACTACAAAGATGACGACGATAAAtaacgcgtctttaccagtttaccacg | |  |  |
| **L2684** | cgccagggttttcccagtcacgtctcggtttcgtcaaacagatgggt | | PCR to amplify upstream fragment of tssM to clone in pRE112 | |
| **L2685** | ttatgttgaaataaccctgaaggcgcattc | |  |  |
| **L2686** | cttcagggttatttcaacataagtgaccattcaactcc | | PCR to amplify downstream fragment of tssM to clone in pRE112 | |
| **L2687** | gaCTCTAGAGGATCCCCGGGTAcgtactggtttgatggccac | |  |  |
| **L758** | TACCCGGGGATCCTCTAGAG | | PCR to amplify the linearized pRE112-lacZα | |
| **L1227** | cgtgactgggaaaaccc | |  |  |
| **L762** | GTTATTGGTGCCCTTAAACG | | To verify insertion of fragment in pRE112 and pRE112-lacZα and used for Sanger sequencing | |
| **L763** | GCATCCAACGCCATTCATGG | |  |  |
| **L1948** | gcttgccagtgaaggcaat | | To verify *slyA* deletion | |
| **L1949** | tcggtaataccgtcggtcgc | |  |  |
| **L2251** | aaatcgatagcgcgttgcag | | To verify *slyA-sfGFP_mcherry* insertion | |
| **L2252** | ccgacacagacacagctataacag | |  |  |
| **L2688** | gtcggactggaagaacaccat | | To verify *tssM* deletion | |
| **L2689** | gcaataaaacccagctggcg | |  |  |
| **L2519** | tgggtggattgcaatcaactctg | | PCR IVA to add mutation GGGA-->CCCA in 5'UTR of *slyA in* pRE112-slyA-sfGFP_RBS-mcherry | |
| **L2520** | gagttgattgcaatccacccaagttgcttagtatgctaactataaggag | |  |  |
| **L2177** | CACCATCACCATCACATCGAAGGGCGCGgatcCggaggaagcggaGAATTGCCCCTAGGATCAG | | PCR to amplify *slyA* optimized from pGEX-6-3P *slyA* to clone in pIBA37 | |
| **L2178** | TTCACAGGTCAAGCTTAGTTAGATTAAGACTGGCTTTCATGCAG | |  |  |
| **L1997** | atcccccgggctgcaggaattcgatatcctggaattcgaattcgccc | | PCR to amplify *slyA* and its promoter to clone in pwsk29-oriT | |
| **L1998** | ggtcgacggtatcgataagcttgatatctgggttaaccgtcatggc | |  |  |
| **L2528** | aagcggtacacctgcgtt | | PCR IVA to add mutation TTTCCC-->TTTGGG in *arcZ* in pwsk29-oriT-*arcZ* (pEGL334) | |
| **L2530** | aacgcaggtgtaccgctttGGGtggtgttggcgcataattcg | |  |  |
| L2693 | ggcctcttcgctattacgccagatcacatcatccggctacg | | PCR to amplify *tssM* and its promoter to clone in pwsk29-oriT | |
| **L2694** | ggtattgacatgttgaaaataatcattacctttttgcg | |  |  |
| L2634 | ctggcgtaatagcgaagagg | | To linearize pwsk29-oriT | |
| **L2635** | ctgcattaatgaatcggccaac | |  |  |
| **L2590** | ctgcaggcatgcaggagg | | To linearize pSEVA421-P*sol*-*gfp-luxCDABE* (pEGL 385) | |
| **L2591** | gagctcgaattcgcgcgg | |  |  |
| **L2718** | GCGGccgcgcgaattcgagctccagactccagcgatcgcac | | To amplify *hcpA* promoter to clone in pSEVA421*-gfp-lux* | |
| **L2719** | TTTTcctcctgcatgcctgcagtgaacggttaggtatgacttcgc | |  |  |
| **L2720** | TTTTcctcctgcatgcctgcagggcccgaaaaaataaggtcg | | To amplify *tssB* promoter to clone in pSEVA421*-gfp-lux* | |
| **L2721** | GCGGccgcgcgaattcgagctcataattcgagttgcaggaaggc | |  |  |
| **L2722** | TTTTcctcctgcatgcctgcagggctccatagctaccgcacacc | | To amplify *vgrGB* promoter to clone in pSEVA421-gfp-lux | |
| **L2723** | GCGGccgcgcgaattcgagctccaggacccgtccaagc | |  |  |
| **L2724** | TTTTcctcctgcatgcctgcagtgaacggttaggtatgacttcgcc | | To amplify *hcpB* promoter to clone in pSEVA421*-gfp-lux* | |
| **L2725** | GCGGccgcgcgaattcgagctccattgtggcacagcgcacatc | |  |  |
| **L2665** | gatgccgttttttcgggagat | | To amplify *hcpA* promoter with 5'-FAM for EMSA | |
| **L2666-FAM** | 5'-FAM-ccttgttgttgaacggttaggtatg | |  |  |
| **L2256-FAM** | 5'-FAM-gaattgtttggcgcggatatg | | To amplify *sol* promoter with 5'-FAM for EMSA | |
| **L2257** | cacaatttacgatataaccagaatgaaaaatc | |  |  |
| **L2669** | tgtttgcgcaatgtttgcaaac | | To amplify *hcpB* promoter with 5'-FAM with L2666-FAM for EMSA | |
| **L2164-FAM** | 5'-FAM-ccggttggttcgtgtatgg | | To amplify T- fragment (part of *slyA* CDS) with 5'-FAM for EMSA | |
| **L2165** | accagcgaaggctgttctat | |  |  |
| **L2109** | tgccgggtaattaacccctt | | RT primer for 5’RACE *slyA* | |
| **L2110** | gggtaattaaccccttgtcttcaagttga | | To amplify the 5’region of *slyA* mRNA from 5’RACE | |
| **L2152-TSO** | CATTGCAAGCAGTGGTATCAAC | |  |  |
| **L707** | agcggataacaatttcacacagga | | To verify insertion of promoter in pSEVA421-gfp-luxCDABE | |
| **L33** | TTAATGCGCCGCTACAGGGCG | |  |  |
