## Supplementary material for "The ArcZ small RNA activates SlyA production to control contact-dependent antibacterial killing and diffusible antifungal activity in *Dickeya solani*": table legends

**Tables S1-S5 legends**

**Table S1: RIL-seq analysis**

A *D. solani* D s0432-1 strain expressing chromosomal Hfq-3×FLAG and an isogenic untagged control strain were grown in LB at 30°C to low cell density (LCD; OD_600_ = 0.5) or high cell density (HCD; OD_600_ = 1.0), with three biological replicates per condition. Hfq-associated RNA‑RNA interactions were identified by RIL-seq and analyzed using ChimericFragments. The Summary LCD and Summary HCD worksheets report global interaction statistics, RNA annotation classes, read distributions, and parameters used for interaction calling. The RIL_SEQ_LCD and RIL_SEQ_HCD worksheets contain the complete interaction datasets for each growth condition, organized according to the identities and orientations of the interacting RNAs. For each RNA pair, the tables report RNA names, RNA classes and functional annotations, number of chimeric reads (nb_ints), number of multimapping reads (nb_multi), number of biological libraries in which the interaction was detected (in_libs), Fisher’s exact test statistics and FDR, base-pairing P value and FDR, interaction-length parameters, and corresponding IPO 2222 and D s0432-1 locus tags. The ArcZ_LCD_focus_analysis and ArcZ_HCD_focus_analysis worksheets extract all interactions involving ArcZ, independently of whether ArcZ was recovered as RNA1 or RNA2, to facilitate analysis of the ArcZ interactome at each cell density. Interactions retained for the analyses presented in the manuscript were supported by ≥5 chimeric reads and a base-pairing FDR ≤ 0.05.

**Table S2: Quantitative proteomic analysis of the Δ*slyA* mutant relative to the WT strain.**

Strains DS753 (Δ*slyA*) and DS49 (WT) were grown in M63 minimal medium supplemented with 1% sucrose at 30°C to OD_600_ = 0.3, with three biological replicates per strain. Proteins were identified and quantified by LC-MS/MS using Proteome Discoverer 3.1 with CHIMERYS and Sequest HT. Protein abundances were normalized to total peptide amount. The raw proteomic data worksheet contains the complete Proteome Discoverer output, including protein accessions, PEP scores, sequence coverage, peptide and PSM counts, and abundance measurements. The analysis worksheet reports differential protein abundance results cross-referenced to *D. solani* IPO 2222 and D s0432-1 locus tags, with log_2_ fold changes (Δ*slyA*/WT) and associated P values. Differentially abundant proteins were defined by a log_2_ fold change ≤ −1 or ≥ 1 and P < 0.05 using the background-based t-test implemented in Proteome Discoverer. A total of 88 proteins showed reduced abundance and 190 showed increased abundance in Δ*slyA* relative to WT.

**Table S3: RNA-seq transcriptomic analysis of the ∆*slyA* mutant relative to the WT strain.**

Strains DS753 (Δ*slyA*) and DS49 (WT) were grown in LB medium at 30°C to OD_600_ = 0.5, with three biological replicates per strain. Total RNA was extracted, depleted of ribosomal RNA, and subjected to paired-end RNA sequencing. Reads were mapped to the *D. solani* IPO 2222 reference genome, and differential gene expression was analyzed using DESeq2 with median-of-ratios normalization. The expression browser worksheet contains raw read counts for each biological replicate. The differential expression worksheet reports results cross-referenced to *D. solani* IPO 2222 and D s0432-1 locus tags, including baseMean, log_2_ fold change (Δ*slyA*/WT), standard error, P-value, and FDR-adjusted P-value (padj) for each gene. Differentially expressed genes were defined by a log_2_ fold change ≤ −1 or ≥ 1 and FDR-adjusted P < 0.05. A total of 175 genes showed reduced expression and 42 showed increased expression in Δ*slyA* relative to WT.

**Table S4:** **Bacterial strains and plasmids used in this study.**

All *Dickeya* solani, Escherichia *coli* and *P. atrosepticum* strains and plasmids used in this study are listed with their genotype, relevant genetic modifications, antibiotic-resistance markers and source or reference.

**Table S5. Oligonucleotides used in this study.**

Sequences are given in the 5′-to-3′ orientation. Lowercase letters denote genomic target-specific regions; uppercase letters denote adapter or universal sequences.
